# Structure-Based Classification of CRISPR/Cas9 Proteins: A Machine Learning Approach to Elucidating Cas9 Allostery

**DOI:** 10.1101/2025.04.05.647395

**Authors:** Sita Sirisha Madugula, Vindi M. Jayasinghe-Arachchige, Charlene R. Norgan Radler, Shouyi Wang, Jin Liu

## Abstract

The CRISPR/Cas9 system is a powerful gene-editing tool. Its specificity and stability rely on complex allosteric regulation. Understanding these allosteric regulations is essential for developing high-fidelity Cas9 variants with reduced off-target effects. Here, we introduce a novel structure-based machine learning (ML) approach to systematically identify long-range allosteric networks in Cas9. Our ML model was trained using all available Cas9 structures, ensuring a comprehensive representation of Cas9’s structural landscape. We then applied this model to *Streptococcus pyogenes* Cas9 (SpCas9) to demonstrate the feature selection process. Using Cα-Cα inter-residue distances, we mapped key allosteric networks and refined them through a two-stage SHAP feature selection (FS) strategy, reducing a vast feature space to 28 critical Lysine-Arginine (Lys-Arg) residue pairs that mediate SpCas9 interdomain communication, stability, and specificity. These Lys-Arg pairs initially shared a 46.5 Å inter-residue distance, but molecular dynamics simulations revealed distinct stabilization behaviors, indicating a hierarchical allosteric network. Further mutational analysis of R78A-K855A (M1) and R765A-K1246A (M2) identified an “electrostatic valley,” a stabilizing network where positively charged residues interact with negatively charged DNA to maintain SpCas9’s structural integrity. Disrupting this valley through direct (M2) or allosteric (M1) mutations destabilized SpCas9’s DNA-bound conformation, leading to distinct pathways for improving SpCas9 specificity. This study provides a new framework for understanding allostery in Cas9, integrating ML-driven structural analysis with MD simulations. By identifying key allosteric residues and introducing the electrostatic valley as a central concept, we offer a rational strategy for engineering high-fidelity Cas9 variants. Beyond Cas9, our approach can be applied to uncover allosteric hotspots in other enzyme regulation and rational protein design.

## Introduction

Clustered Regularly Interspaced Short Palindromic Repeats and CRISPR-associated protein 9 (CRISPR/Cas9) is a revolutionary genome editing technology that allows precise targeted changes to the DNA with unprecedented ease^1–3^. Originally discovered as a natural defense mechanism in bacteria to cut the DNA of invading viruses, CRISPR/Cas9 has since been adapted as a gene-editing technology^4–6^. CRISPR/Cas9 offers numerous advantages, allowing for precise DNA modifications such as gene knockout, introducing specific mutations, or inserting new genetic material. This precision has profound implications in medical research, agriculture, biotechnology and many more fields^7–10^. CRISPR systems are broadly classified into two classes. Class 1 systems, which include types I, III, and IV, utilize multi-protein complexes for their function. In contrast, Class 2 systems, which include types II, V, and VI, rely on single effector proteins, with Cas9 being the most well-studied among them^11–13^. The Type II CRISPR/Cas9 system has gained prominence due to its simplicity and efficiency, making it a versatile tool for genome editing in various organisms^6,14–16^.

Central to the CRISPR/Cas9 system is Cas9, an RNA-guided nuclease that can be programmed to cleave a specific DNA or RNA sequence using a suitable guide RNA (sgRNA). *Streptococcus pyogenes* Cas9 (SpCas9) is one of the most well-studied Cas proteins, with versatile applications in therapeutic and applied fields owing to its simple design and versatility^17,18^. Despite the numerous advantages of Cas9, it suffers from inherent limitations like specificity and off-target effects, wherein the enzyme cuts DNA at unintended sites^19,20^. This lack of specificity leads to unwanted cleavage, having implications for therapeutic applications of CRISPR/Cas9 systems; particularly for genetic diseases where precise gene editing is paramount. Addressing this challenge requires a deeper understanding of the molecular mechanisms governing Cas9 function-particularly the role of allosteric regulation in modulating its catalytic activity and DNA binding properties.

Allostery, the process by which conformational changes at one site in a protein influence distant functional sites, is central to Cas9 activity. Considerable efforts to enhance the technology are centered on refining the editing properties of the Cas9 protein^21–24^. Many computational and experimental studies have been conducted to understand the catalytic and allosteric mechanisms in Cas9^25–29^. Upon recognizing the protospacer adjacent motif (PAM), Cas9 undergoes a series of conformational transitions that facilitate DNA unwinding, guide RNA (sgRNA) hybridization, and subsequent cleavage of the target strand. These transitions are orchestrated through allosteric communication between key structural domains, including the HNH, RuvC, REC, and Bridge Helix (BH) domains. Molecular Dynamics (MD) and Cas9 experimental studies reveal that the REC1-3 domains are crucial in DNA recognition. These domains undergo a conformational change, facilitating a large transition of the catalytic HNH domain towards the DNA cleavage site, effectively locking the HNH domain in place and enabling precise DNA cleavage^30–32^. Therefore, the specificity of the Cas9 nuclease is heavily dependent on the REC-HNH-RuvC allosteric communication^27,27,29,33,34^. Exploiting the allosteric signaling in Cas9, many high-fidelity SpCas9 variants are designed by rationally mutating residues in various critical domains like REC, RuvC, HNH, PI and BH to achieve improved specificity and reduced off-target effects compared to wild-type *Sp*Cas9. Many MD, QM/QM, and FRET studies have explored the mechanism of specificity improvement in Cas9 variants. They involve strategies like (a) enhanced proofreading that allows effective discrimination between on-target and off-target sequences, (b) reducing non-specific interactions by selectively disrupting electrostatic or hydrophobic interactions between Cas9 and off-target DNA sequences, (c) conformational changes in Cas9 structure that alter the accessibility of the target DNA thereby influencing the stability of Cas9-DNA complex^35–37^. While these studies focus on the mechanism through which these mutations influence Cas9 specificity and off-target effects, they do not explicitly address how the long-range communications between residue pairs within Cas9 structure influence its overall stability and functional integrity. The precise nature of long-range communication and the key residue pairs mediating them remain poorly understood.

To systematically investigate allosteric networks within Cas9, we employ a structure-based machine learning (ML) approach that encodes inter-residue distances as predictive features. By analyzing a dataset of Cas9 and non-Cas9 proteins, we identify Lysine-Arginine (Lys-Arg) interactions spanning 5-75Å as critical mediators of allosteric signaling. These interactions bridge functional domains, playing a pivotal role in maintaining Cas9 stability and regulating its specificity. Mutation studies further reveal that disrupting key Lys-Arg pairs (e.g., R78-K855 in BH-HNH of SpCas9) leads to widespread conformational changes, highlighting their role in long-range allosteric communication. By integrating ML-driven structural analysis with molecular dynamics (MD) simulations, this study provides a novel framework for understanding Cas9 allostery. The identification of allosteric residue pairs offers a rational basis for engineering Cas9 variants with enhanced specificity, reduced off-target effects, and improved therapeutic potential.

## 2. Materials and Methods

### 2.1. Datasets

#### Cas9 dataset

All “CRISPR/Cas9 endonuclease” protein structures available in the Research Collaboratory for Structural Bioinformatics Protein Data Bank (RCSB-PDB)^38^ were manually downloaded, and low-resolution and single-domain structures were discarded. Thereafter, 53 remaining CRISPR/Cas9 structures containing all characteristic domains of the Cas9 protein, are carefully inspected and included in the dataset. Subsequently, all Cas9 crystal structures are preprocessed to retain only one of the chains and delete all heteroatoms. Similarly, protein structures from the AlphaFold2 database (2023)^39,40^ with the keyword “CRISPR/Cas9 endonuclease” which at the time of collection included 2377 structures are manually retrieved, and incomplete or single-domain structures are excluded.

#### Non-Cas9 dataset

All non-Cas9 structures are also collected from the RCSB-PDB database^38^, comprising of oxidoreductases, transferases, hydrolases, lyases, isomerases, ligases, and translocases. To ensure effective classification, we selected proteins within a range of 700–2100 amino acids length matching the approximate size range of Cas proteins.

### 2.2. Feature generation: C_-α_ to C_-α_ distance matrix generation

The alpha Carbon (Cα)-(Cα) distances between all amino acid residue pairs within the 3D structures of Cas9 and non-Cas9 proteins were calculated using the BioPython package^41^. The distance matrix obtained is a representation of the protein backbone.

**Table 1.** Datasets used in the study.

| S. No. | Source Database | No. of structures |
| --- | --- | --- |
| Cas9 | RCSB-PDB | 53 |
|  | AlphaFold-2 | 982 |
| Non-Cas | RCSB-PDB | 1033 |
| TOTAL |  | 2068 |

### 2.3. Feature Processing: Lower diagonal distance processing

#### 2.3.1. Lower diagonal flattening of the distance matrix

A distance matrix is a square matrix with identical upper and lower diagonals. To reduce computational cost, only the lower diagonal of the distance matrices was used. The lower diagonal is flattened into five columns, namely, residue pair number, distances, residue pair type, merged residue pair type, and protein. Table 2 shows a representation of the flattened lower diagonal using the structure with PDB ID 4CMP as an example. The column “residue pair number” represents the exact pair of residues in the protein structure, the “distance” column represents the distance between the said residue pair number, and the “residue pair type” column represents the base name of the said residue pair number. The “merged residue pair type” column gives a uniform name for a residue-pair type and its converse. This column ensures that the residue pair type number and its converse are both included under the same base name. The “protein” column represents the PDB ID of the protein.

**Table 2.** Flattening lower diagonal of distance matrix for PDB ID 4CMP.

| Residue Pair number | Distance | Residue pair type | Merged residue pair type | Protein |
| --- | --- | --- | --- | --- |
| ALA1023-ALA1053 | 19.354 | ALA-ALA | ALA-ALA | 4CMP |
| ALA50-ALA292 | 32.941 | ALA-ALA | ALA-ALA | 4CMP |
| ALA50-ALA287 | 25.450 | ALA-ALA | ALA-ALA | 4CMP |
| ALA50-ALA280 | 22.454 | ALA-ALA | ALA-ALA | 4CMP |
| ALA1081-ALA1140 | 47.155 | ALA-ALA | ALA-ALA | 4CMP |
| ... | ... | ... | ... | ... |
| ... | ... | ... | ... | ... |
| ... | ... | ... | ... | ... |
| ALA50-ALA1306 | 26.678 | ALA-ALA | ALA-ALA | 4CMP |
| ALA243-GLY660 | 55.513 | ALA-GLY | ALA-GLY | 4CMP |
| ALA243-GLY865 | 44.600 | ALA-GLY | ALA-GLY | 4CMP |
| ALA91-GLY736 | 48.589 | ALA-GLY | ALA-GLY | 4CMP |
| GLY8-ALA725 | 23.900 | GLY-ALA | ALA-GLY | 4CMP |
| GLY358-ALA1053 | 72.336 | GLY-ALA | ALA-GLY | 4CMP |
| GLY358-ALA1320 | 59.824 | GLY-ALA | ALA-GLY | 4CMP |

#### 2.4.2. Distance table generation

In this step, the flattened table from the previous step is transformed into a 210-column distance table (Table 3), where each column corresponds to one of the 210 possible residue pair types among the 20 known amino acids. Using the “merged residue pair type” column from the flattened table, distances for each residue pair type are segregated into individual columns. Each column represents all Cα-Cα distances between residues of a given pair type across the entire protein structure, resulting in a 2D distance table. Table 3 provides a schematic representation of this distance table using PDB ID 4CMP as an example.

**Table 3.**
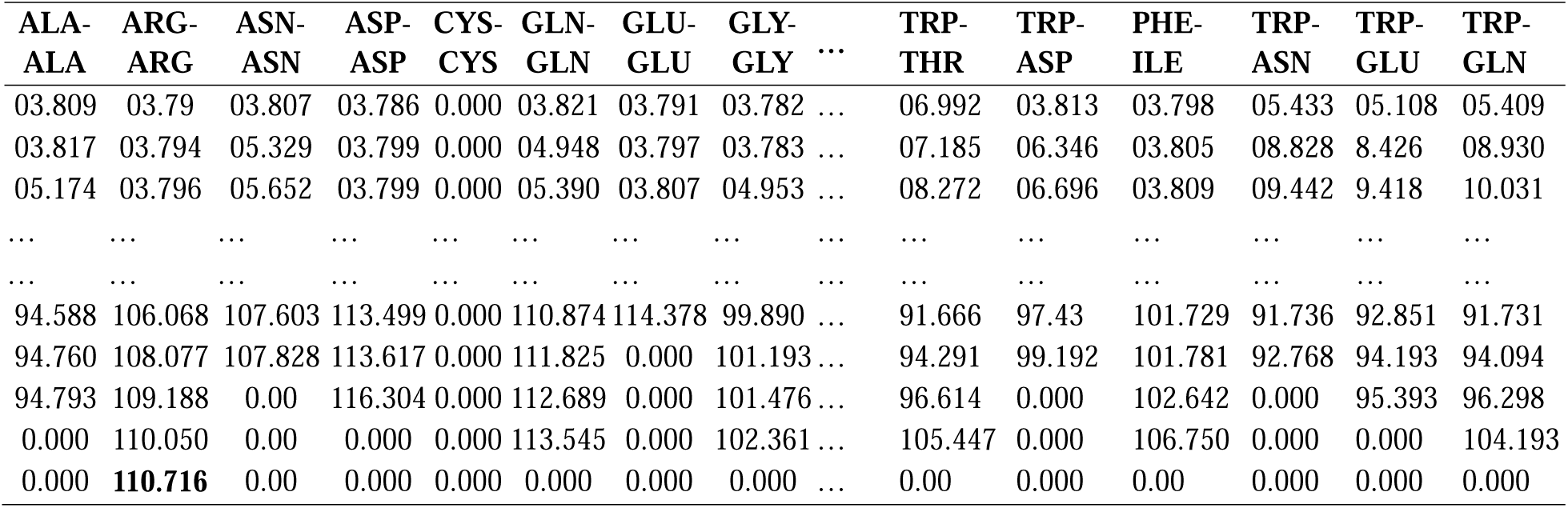
A schematic representation of the distance table for PDB ID 4CMP.

| ALA-ALA | ARG-ARG | ASN-ASN | ASP-ASP | CYS-CYS | GLN-GLN | GLU-GLU | GLY-GLY | ... | TRP-THR | TRP-ASP | PHE-ILE | TRP-ASN | TRP-GLU | TRP-GLN |
| --- | --- | --- | --- | --- | --- | --- | --- | --- | --- | --- | --- | --- | --- | --- |
| 03.809 | 03.79 | 03.807 | 03.786 | 0.000 | 03.821 | 03.791 | 03.782 | ... | 06.992 | 03.813 | 03.798 | 05.433 | 05.108 | 05.409 |
| 03.817 | 03.794 | 05.329 | 03.799 | 0.000 | 04.948 | 03.797 | 03.783 | ... | 07.185 | 06.346 | 03.805 | 08.828 | 8.426 | 08.930 |
| 05.174 | 03.796 | 05.652 | 03.799 | 0.000 | 05.390 | 03.807 | 04.953 | ... | 08.272 | 06.696 | 03.809 | 09.442 | 9.418 | 10.031 |
| ... | ... | ... | ... | ... | ... | ... | ... | ... | ... | ... | ... | ... | ... | ... |
| ... | ... | ... | ... | ... | ... | ... | ... | ... | ... | ... | ... | ... | ... | ... |
| 94.588 | 106.068 | 107.603 | 113.499 | 0.000 | 110.874 | 114.378 | 99.890 | ... | 91.666 | 97.43 | 101.729 | 91.736 | 92.851 | 91.731 |
| 94.760 | 108.077 | 107.828 | 113.617 | 0.000 | 111.825 | 0.000 | 101.193 | ... | 94.291 | 99.192 | 101.781 | 92.768 | 94.193 | 94.094 |
| 94.793 | 109.188 | 0.00 | 116.304 | 0.000 | 112.689 | 0.000 | 101.476 | ... | 96.614 | 0.000 | 102.642 | 0.000 | 95.393 | 96.298 |
| 0.000 | 110.050 | 0.00 | 0.000 | 0.000 | 113.545 | 0.000 | 102.361 | ... | 105.447 | 0.000 | 106.750 | 0.000 | 0.000 | 104.193 |
| 0.000 | **110.716** | 0.00 | 0.000 | 0.000 | 0.000 | 0.000 | 0.000 | ... | 0.00 | 0.000 | 0.00 | 0.000 | 0.000 | 0.000 |

In the distance table, each column signifies a specific residue pair within the protein. Within each column, the number of distances corresponds to the count of Cα-Cα distances of the said residue pair type within the entire protein. Since a protein can have different numbers of each residue pair type, different residue pair types exhibit varying counts of distances in their respective columns. Therefore, the distance table has a varying number of rows within its 210 columns. To process these irregular distance tables using Python code, it becomes imperative to ensure uniform dimensions across all columns. To achieve this, a systematic approach is adopted: first, we identified the column with the highest count of distances, which serves as a reference (boldfaced in the Arg-Arg column of Table 3). Next, leveraging the highest count from this reference column, we pad the remaining cells of all other columns with zeros until they match the highest count in the reference column. This procedure ensures consistent dimensions of the distance table for a protein, facilitating its easy processing in the next steps. While this procedure ensures uniform dimensions of distance table for a protein, it is important to note that the overall dimensions of distance tables can still vary among different proteins. This variability arises due to the differing sizes of the proteins leading to different numbers of residues in each protein structure. Consequently, proteins with a higher residue count have distance tables with larger dimensions, while proteins with slightly fewer residues have smaller distance tables. This inherent variation in dimensionality reflects the unique structural characteristics of each protein and underscores the importance of accounting for protein-specific attributes when analyzing distance data.

#### 2.4.3. Transformation into histogram bins

Since different proteins have different dimensions of the distance table, it is necessary to convert them into a uniform dimension that comprehends the unique distances of Cas9 and non-Cas9 proteins of the dataset. To ensure comparability across different proteins we employed a histogram transformation approach. This technique grouped the distances from the distance tables into uniform histogram bins. In this study, since we performed two rounds of feature selection (FS), we conducted the histogram transformation process twice, employing two different bin intervals. We first computed the global minimum and maximum distance values (0.3 and 165Å, respectively) across all Cas9 and non-Cas9 proteins of the dataset. Based on these values, we first employed a broad bin interval of 5Å, which resulted in 34 bins, from which 33 bins capture all distances within our dataset proteins, with an additional bin accounting for zeros added for padding the distance tables in the previous step. This broad binning approach is intended to get an overall idea of the most important residue types and their inter-residue distances across the entire Cas structure. In the second round, we refined the selection process by focusing on Lysine-Arginine pairs within the 35-65Å distance range that correspond to the top 6 bits of the first round of FS. We employed a second histogram transformation, this time using a smaller bin interval of 0.1Å. This finer granularity led to a total of 304 bins, where 300 bins cover all 35-65Å distances, and three additional bins account for padded zeros and one each for distances below 35Å and above 65Å, respectively. This granularity allows for a more precise investigation of specific residue-pair distances influencing the Cas9 protein structure. Tables 4a and 4b represent the 33 and 300 columns obtained, respectively, in the first and second rounds of FS for the structure with the PDB ID 4CMP.

**Table 4a.**
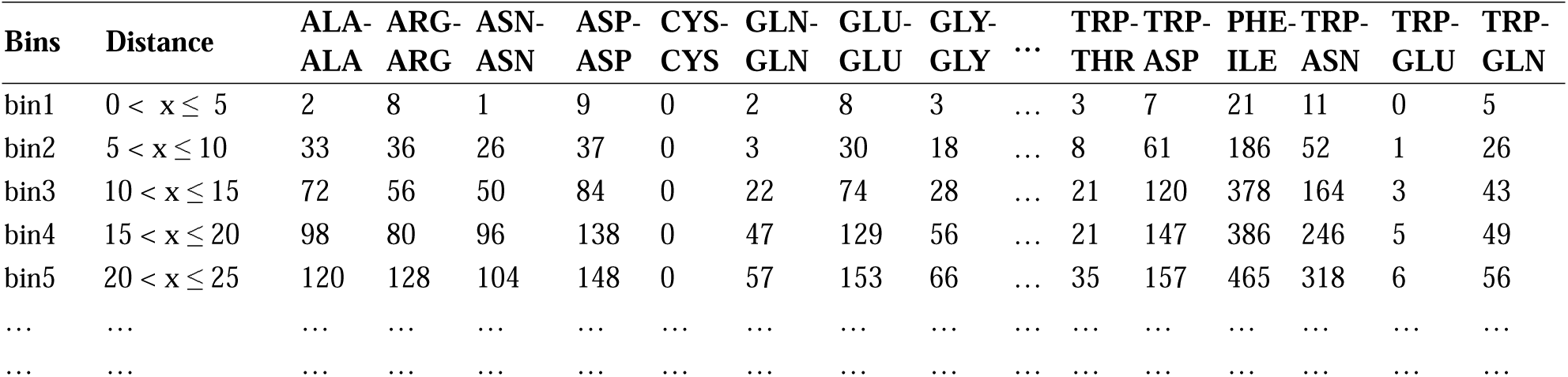
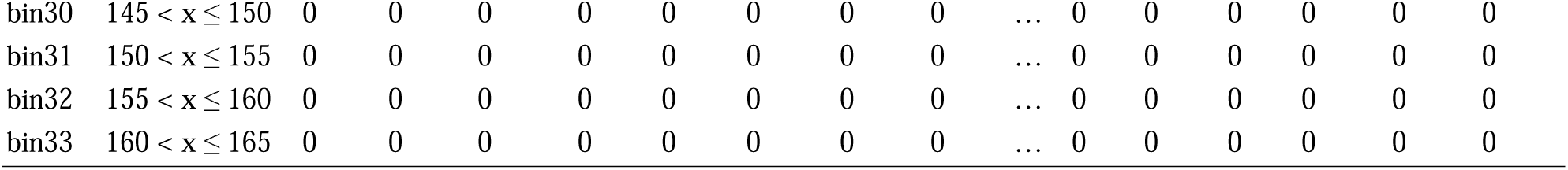
A schematic representation of the 33 histogram bins for PDB ID 4CMP.

**Table 4b.**
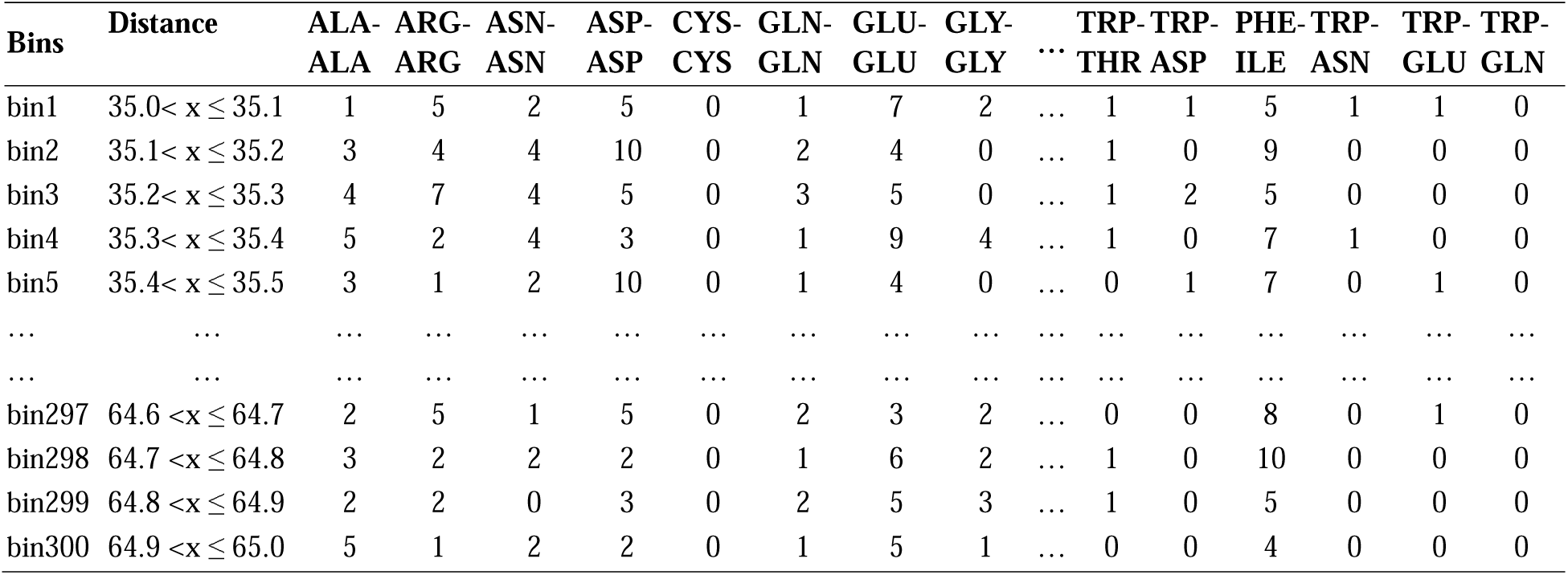
A schematic representation of the 300 histogram bins for PDB ID 4CMP.

#### 2.4.4. Concatenated vector generation

Next, the 300 bin values for each residue pair in Table 4b are concatenated vertically, i.e. bin-wise, to generate a 1D vector of size 63000 bits corresponding to bin values of all 210 residue pairs in the table. In the first round of histogram transformation, the same procedure produced a 6930-bit vector after concatenating the 33 bin values for all 210 columns shown in Table 4a.

### 2.5. Two-Stage Feature Selection

The 63000-bit vector is generated for all proteins of the dataset, forming a 63000 X 2068 data frame on which we performed Shapley Additive exPlanations (SHAP) feature selection. SHAP is a game theory-based feature selection method that determines the relative feature importance of each feature towards the model performance^42,43^. Using the TreeSHAP algorithm, we calculated the Mean Absolute SHAP values, which represent the global influence of each feature on model predictions^44^. Unlike traditional importance-based methods, SHAP accounts for feature interdependencies while ranking their contributions to the final prediction, offering robust and interpretable results. In our study, rather than using raw distances, we transformed residue-pair distances into vector bits, where each bit encodes distances within a specific bin range. Two rounds of SHAP feature selection are conducted with distinct objectives. In the first round, we performed SHAP FS on the 6390-bit concatenated vectors covering all distance ranges of dataset proteins to initially assess their relevance to Cas9 stability. To minimize variance, we averaged SHAP importance scores across 15 iterations of feature selection. Building on the insights of the first round, we narrowed our search to a smaller 35-65Å distance range and performed SHAP on 63,000-bit vector to determine specific residue-pair distances critical for the Cas9 structure. This identified many important residue-pair numbers and their distances, which are important for the Cas9 structure.

### 2.6. Random Forest ML model building

To optimize model performance, the top 6 Lysine-Arginine bits identified from the first round of SHAP are used to build RF classification models to differentiate Cas9 from non-Cas9 proteins. The top 6 bits are chosen based on their Mean Absolute SHAP values since it is observed that these are the minimum number of bits that resulted in significant performance improvements. Similarly, in the second round of feature selection, we used the top 2 bits for RF model building.

### 2.7. Evaluation metrics

We evaluated our models using six evaluation metrics: accuracy, recall, F1 score, precision, specificity, and Area Under the Receiver Operating Characteristic Curve (AUC-ROC) on the test dataset.

### 2.8. Experimental Settings

Here we classified Cas9 proteins versus non-Cas9 proteins using a novel, data-driven approach that encodes the 3D structural information of Cas9 proteins into uniform bit representations. This preprocessing method involves converting the backbone of Cas9 protein structure into a Cα distance matrix, which is subsequently transformed into concatenated bits and used for classification. The dataset is split into training and test sets in 80:20 ratio using a stratified split to ensure balanced classes. SHAP feature selection is then applied to the concatenated vector to identify important bits. The top 13 bits from the first round of feature selection are used further for model building and test set prediction. RF is employed for model building, with hyperparameter tuning conducted using Scikit-Learn’s GridSearchCV with 5-fold cross-validation. We tuned 5 parameters ‘n_estimators,’ ‘max_depth,’ ‘min_samples_leaf,’ ‘min_samples_split,’ and ‘max_features’ for this study. Since the Cas9 vs. non-Cas9 classification task is a binary problem, accuracy is used for evaluation. To ensure robustness and cover model variance, the entire experiment was repeated 15 times. In the second round of FS, the same procedure is applied, but with a narrower distance range of 35-65Å, a smaller bin interval of 0.1Å, and a larger vector size, following which RF models are built using the top 2 bits. This focused approach enabled us to zoom in on key residue pairs critical for maintaining Cas9 structure stability and functional integrity. In both rounds, the number of bits selected for building RF models is based on the Mean Absolute SHAP values of the top bits identified in each round of FS. The objective is to identify the minimum number of bits that still resulted in a significant performance improvement. This led to selecting 6 bits in the first round and 2 bits in the second round.

### 2.9. Molecular dynamics (MD) simulations

**Structural Model:** To reconstruct the active-state cryo-EM structure of wild-type (WT) SpCas9 (PDB ID: 6O0Y^45^) and address missing residues (175–310, 713–717, 1002–1075), nucleotides, and metal ions, we employed a stepwise approach. Using a previously simulated HNH precatalytic structure based on the X-ray structure of Cas9 (PDB ID: 5F9R^46^), missing nt-DNA was superimposed and manually completed. To maintain the conformation of the RuvC catalytic center, key regions were replaced with corresponding residues from 6O0Y. Mg² ions in the HNH center were added from prior simulations, while their positions in the RuvC domain were modeled using the Mn²-bound structure (PDB ID: 4CMQ^47^). This hybrid model integrates experimental data and manual refinements for structural and functional analyses. Furthermore, the two mutated systems M1(R78A-K855A) and M2(R765A-K1246A) were constructed by mutating corresponding Lysine and Arginine residues in each pair to Alanine, respectively.

**MD setup:** The molecular dynamics (MD) simulations were performed using AMBER18^48^. The LEaP ^49^module was used to add hydrogen atoms, neutralize the system with counterions, and solvate the structure in a TIP3P^50^ water box extending 12 Å from the complex surface. The ff14SB^51^, OL15^52^, and OL3^53^ force fields were employed for proteins, DNA, and RNA, respectively. Systems were minimized for 10,000 cycles using a combination of the steepest descent (1,000 cycles) and conjugate gradient (9,000 cycles) algorithms, with restraints on heavy atoms. Heating to 310 K was performed using Langevin dynamics^54–56^ (collision frequency: 2 ps ¹), followed by a 1 ns equilibration in the NPT ensemble with reduced heavy-atom restraints. Production simulations were run without restraints in the NPT ensemble, using SHAKE^57^ to constrain bonds involving hydrogens and smooth particle mesh Ewald^58^ for long-range Coulomb interactions, with a 10 Å cutoff for non-bonded interactions. Duplicate simulations for all 3 systems (WT, M1, and M2) with of at least 300 ns each were conducted with a 2 fs time step, saving trajectories every 2 ps.

**Structural analysis:** The CPPTRAJ^59^ module in AMBER was used to perform root mean square deviation (RMSD) calculations, distance analysis, and clustering. Additionally, the molecular mechanics/generalized Born surface area (MM/GBSA) ^60–62^ method was employed to calculate the binding enthalpies of the WT, M1, and M2 systems, considering nt-DNA as the ligand and Cas9 as the receptor.

## 3. Results and Discussion

### 3.1 A Machine Learning model to identify critical residue-pairs for Cas9

Understanding the allosteric regulation of Cas9 is critical for optimizing its function. Cas9 undergoes a series of conformational transitions during DNA recognition and cleavage, facilitated by long-range allosteric communication between functional domains, including REC, HNH, RuvC, and the Bridge Helix (BH). While previous studies have identified key structural rearrangements in Cas9, the specific residue pairs mediating these allosteric transitions remain largely unknown. To systematically identify these allosteric networks, we developed a structure-based machine learning (ML) model that leverages inter-residue distances as predictive features. We employ a unique structure-based approach leveraging Cα distances within Cas9 structure as features to train Random Forest (RF) models. This strategy distinguishes Cas9 proteins from non-Cas9 proteins while simultaneously identifying residue pairs and distances critical for the overall stability and functional integrity of Cas9 structure. By utilizing 3D structural features for classification, we effectively embed spatial arrangements of amino acids and key stability factors which sequence-based models might overlook.

**Figure 1.**
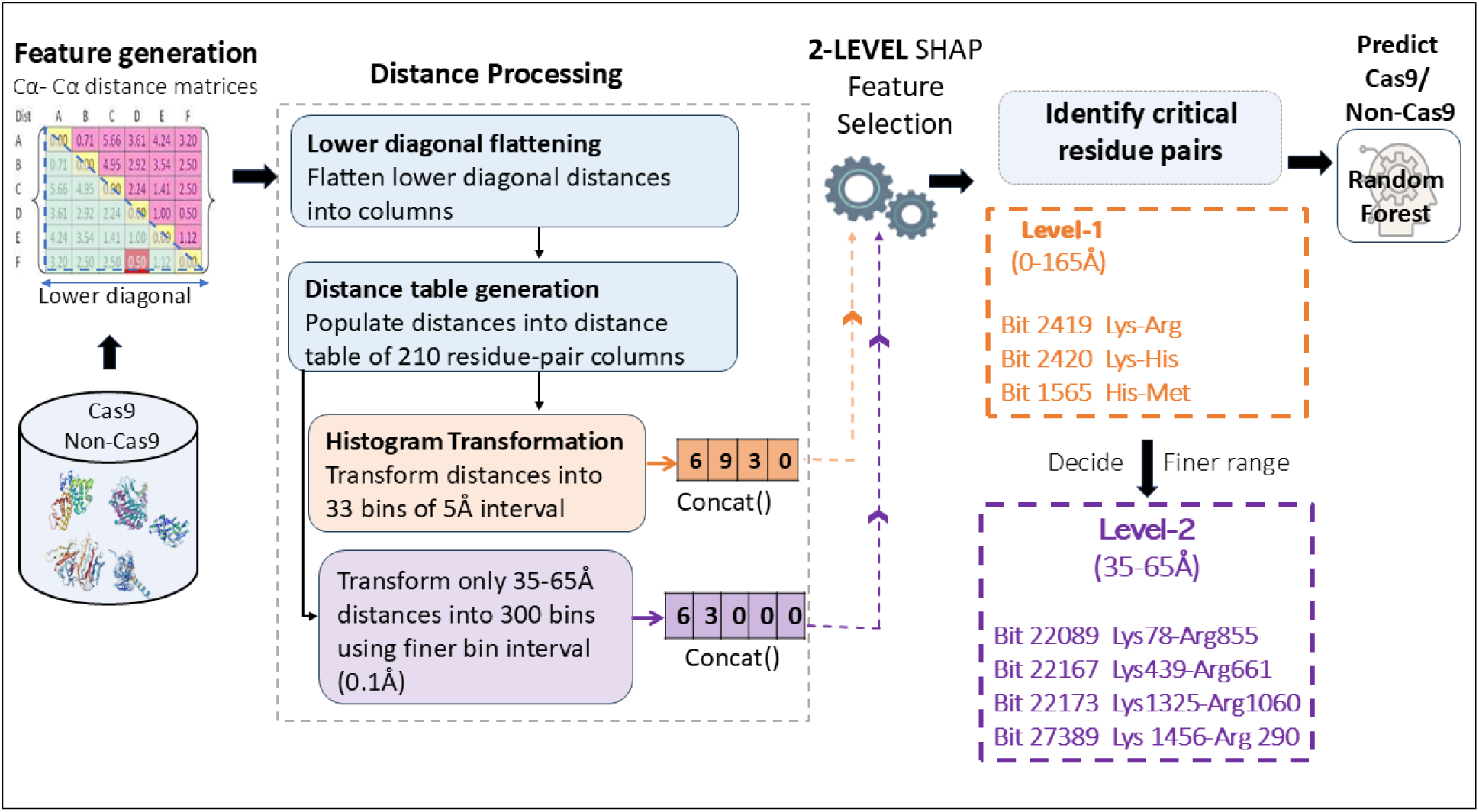
The overall workflow of the machine learning model. All Cα-to-Cα inter-residue distances within Cas9 and non-Cas9 protein structures are first converted into distance matrices. Lower diagonal distances are then flattened into columns of residue-pair and distances. The distances are then sorted into a distance table of 210 columns, representing all possible pairs between the 20 natural amino acids. The distances are further transformed into histograms in two stages: a course (broad range) 5Å binning in the first round and a finer 1Å binning in the second. Bin-wise concatenation of values generated 1D vectors of 6930 and 63,000 bits in the first and second rounds respectively. This is followed by a two-stage feature selection on the two different vectors using the SHAP FS method. The first level FS aims to identify the most important type of residue-pair and range across all proteins of the dataset. A finer distance range is selected based upon the top residue-type identified in level 1 and a second round of FS is carried out within this distance range to refine the results using the 63,000-bit vector. The top bits identified in both levels are used to develop respective Random Forest models that can distinguish Cas9 from Non-Cas9 proteins.

We took the top 13 bits of the first round FS and built RF models for Cas9 vs. non-Cas9 classification. To refine the results further, we focused on the distance range of the top 6 bits (35-65Å) and performed a second round of FS. Subsequently the top 2 bits from the second round are used further to develop Cas9 vs. non-Cas9 RF classification models. The test set performance of RF models built using the top 13 and top 2 bits identified in the first and second rounds of FS are shown in Tables 5 and 6 respectively.

**Table 5.** Test set performance of RF model built using top 13 bits of first round averaged over 15 runs.

| S. No. | Metrics | Values |
| --- | --- | --- |
| 1 | Accuracy | 0.979±0.006 |
| 2 | Specificity | 0.977±0.009 |
| 3 | Precision | 0.977±0.008 |
| 4 | Recall | 0.981±0.009 |
| 5 | F1-score | 0.979±0.006 |

As seen in Tables 5 and 6, the models developed using the top 6 and top 2 bits from the first and second round FS, respectively, performed well across all evaluation metrics. They achieved a high average accuracy of up to 97.0%, along with similar performance in Specificity, Precision, Recall, and F1-score. This shows that among the 210 pairs, Lysine-Arginine residue-pair types, particularly in the 5-75Å distance range, are the most important in distinguishing Cas9 from non-Cas9 proteins.

Interestingly, our second-stage FS models (Table 6), built using fewer Lysine-Arginine bits, showed nearly close performance to the first-stage models developed using 13 bits, demonstrating the effectiveness of our two-stage FS approach in identifying the most important residue pairs governing Cas9 structure with fewer features. The approach effectively reduces model complexity while maintaining accuracy, underscoring the importance of Lysine-Arginine pairs in Cas9 stability. The method also enables identifying other residue types after Lys-Arg that can be important for the cas9 structure identified based on their SHAP feature importance. Additionally, the method revealed key structural insights, which are difficult to capture using sequence-based methods alone, holding potential applications in other protein studies.

**Table 6.** Test set performance of RF model using top 2 bits of second round averaged over 15 runs.

| S. No. | Metrics | Values |
| --- | --- | --- |
| 1 | Accuracy | 0.968±0.008 |
| 2 | Specificity | 0.967±0.010 |
| 3 | Precision | 0.967±0.010 |
| 4 | Recall | 0.969±0.011 |
| 5 | F1-score | 0.968±0.008 |

### 3.2. Long-range Lys-Arg pairs are critical to distinguish Cas9 from non-Cas9 proteins

Since a broad bin interval 5Å is used in the first round FS, every bit identified by SHAP contains many residue pairs within each bit. Table 7 shows the top 13 bits from the first round FS. For other bits, their importance and ranks are given in **SI.**

**Table 7.** Top 13 bits obtained from the first round of feature selection.

| S. No. | Bits | Residue-pair type | Distance range (Å) | Average SHAP value |
| --- | --- | --- | --- | --- |
| 1 | bit2419 | Lysine-Arginine | **45-50** | 0.010064 |
| 2 | bit2420 | Lysine-Arginine | **50-55** | 0.009805 |
| 3 | bit2418 | Lysine-Arginine | **40-45** | 0.009627 |
| 4 | bit2421 | Lysine-Arginine | **55-60** | 0.009531 |
| 5 | bit2422 | Lysine-Arginine | **60-65** | 0.009386 |
| 6 | bit2417 | Lysine-Arginine | **35-40** | 0.008525 |
| 7 | bit2416 | Lysine-Arginine | **30-35** | 0.007522 |
| 8 | bit2423 | Lysine-Arginine | **65-70** | 0.007154 |
| 9 | bit2411 | Lysine-Arginine | **05-10** | 0.006852 |
| 10 | bit2415 | Lysine-Arginine | **25-30** | 0.006517 |
| 11 | bit2413 | Lysine-Arginine | **15-20** | 0.006451 |
| 12 | bit2414 | Lysine-Arginine | **20-25** | 0.006402 |
| 13 | bit2424 | Lysine-Arginine | **70-75** | 0.005481 |

As seen in Table 7, 13 of the top bits are all Lysine-Arginine pair type within the 5-75Å distance range, demonstrating that Lysine-Arginine is the most important pair-type for the overall structure of Cas9 protein. This result coincides with the fact that lysine and arginine are the most associated residues with critical Cas9 activities like DNA binding, stabilization of RNA-DNA heteroduplex, allosteric communication, and DNA cleavage. These 13 bits correspond to short- and long-range distances over a wide range of 5Å to 75Å (boldfaced). While short-range distances within proteins stabilize local structural elements, long-range distances determine the overall integrity and functional activities of proteins, suggesting their role in both local structural stabilization and long-distance allosteric communication.

Further, we analyzed the top 13 bits of three well-studied bacterial species, namely *Streptococcus pyogenes* (SpCas9)*, Campylobacter jejuni* (CjCas9) and *Neisseria meningitidis* (NmeCas9). We compared the number of lysine-arginine residue pair numbers in top 13 bits of all three bacterial Cas9 species shown in Figure 2.

**Figure 2.**
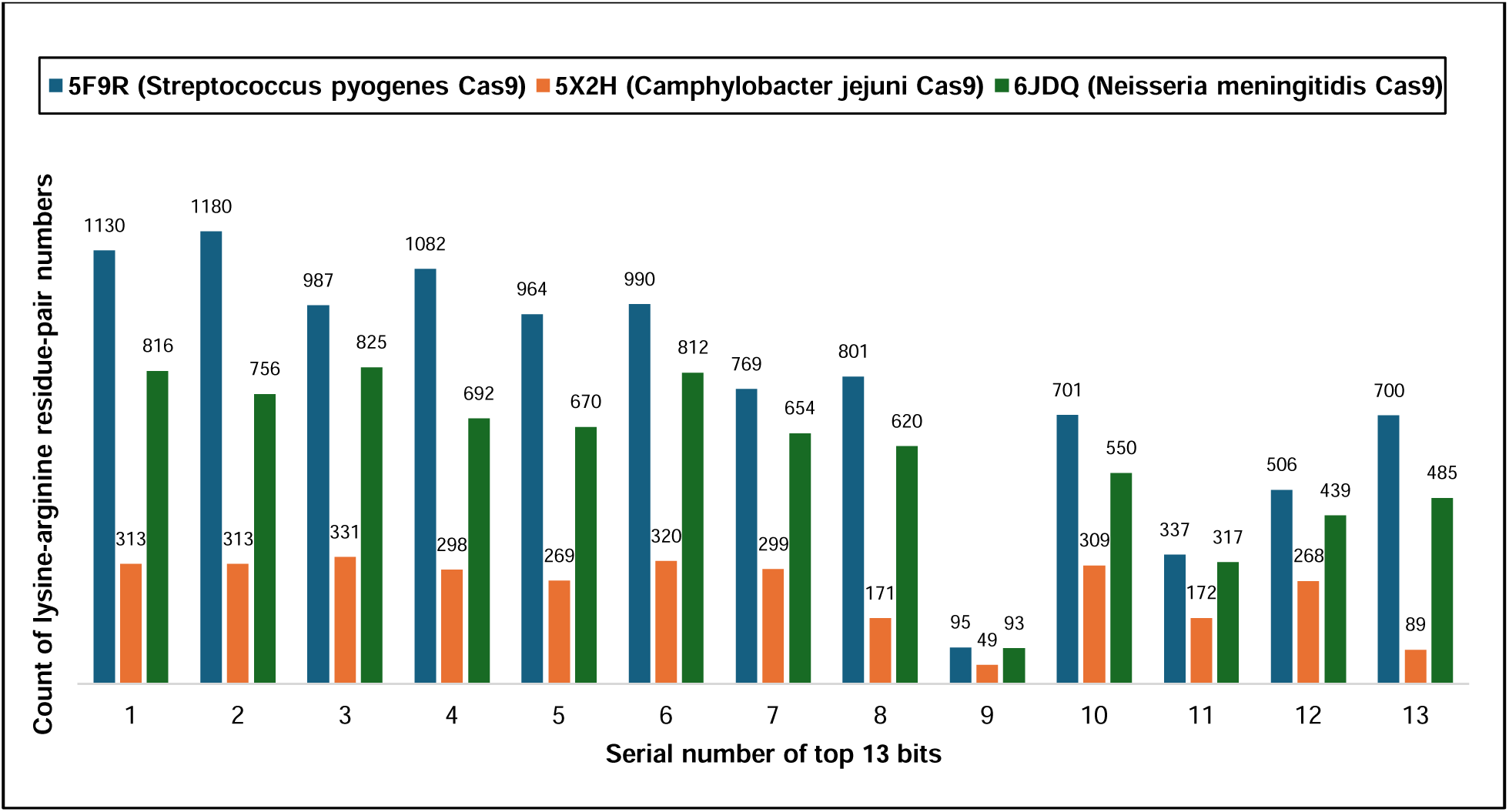
Comparative analysis of the top 13 bits among three well-studied Cas9 species. Comparative count of Lysine-Arginine residue-pair numbers within the top 13 bits in the three bacterial Cas9 structures.

The analysis revealed a high total count of Lysine-Arginine residue pairs in these bits, with 10,242 pairs in SpCas9, 3201 pairs in CjCas9, and 7729 pairs in NmeCas9, as shown in Figure 2. The 10,242 residue pairs in SpCas9 are given in supporting Information (SI). We noticed that the residue-pair counts in a certain distance range, such as Bit 9 (5-10 Å), are much smaller than other bits. Therefore, to refine the analysis, we focused on the top 6-bit range spanning in 35-65Å (Table 7) and performed a second round of SHAP feature selection. Since SpCas9 is the most extensively utilized Cas9 variant in experimental efforts to improve editing properties, such as specificity and off-target effects, our forthcoming sections will center on the analysis of critical residue pairs within SpCas9.

### 3.3. Identification of 28 Lys-Arg pairs as the allosteric nodes for SpCas9

Following the initial identification of Lys-Arg interactions as dominant allosteric mediators, we refined our analysis using a second round of feature selection within 35-65Å, using higher resolution binning (0.1Å intervals). Using a finer bin interval of 0.1 Å and grouping distances into 300 bins increased the bin count from 33 to 300 but reduced the number of lysine-arginine residue pairs per bin significantly, thus reducing the number of lysine-arginine distances to be analyzed within each bit, further helping in quickly identifying the important pairs. The finer binning enabled more refined and faster identification of the most crucial residue pairs, streamlining the feature selection process. The top bit of SpCas9 obtained from the second round of FS identified 28 lysine-arginine pairs within 46.5Å-46.6Å distance range that have significance with respect to the SpCas9 stability, and possibly other properties like specificity and off-target activity. These residue pairs are shown in Table 8.

**Table 8.**
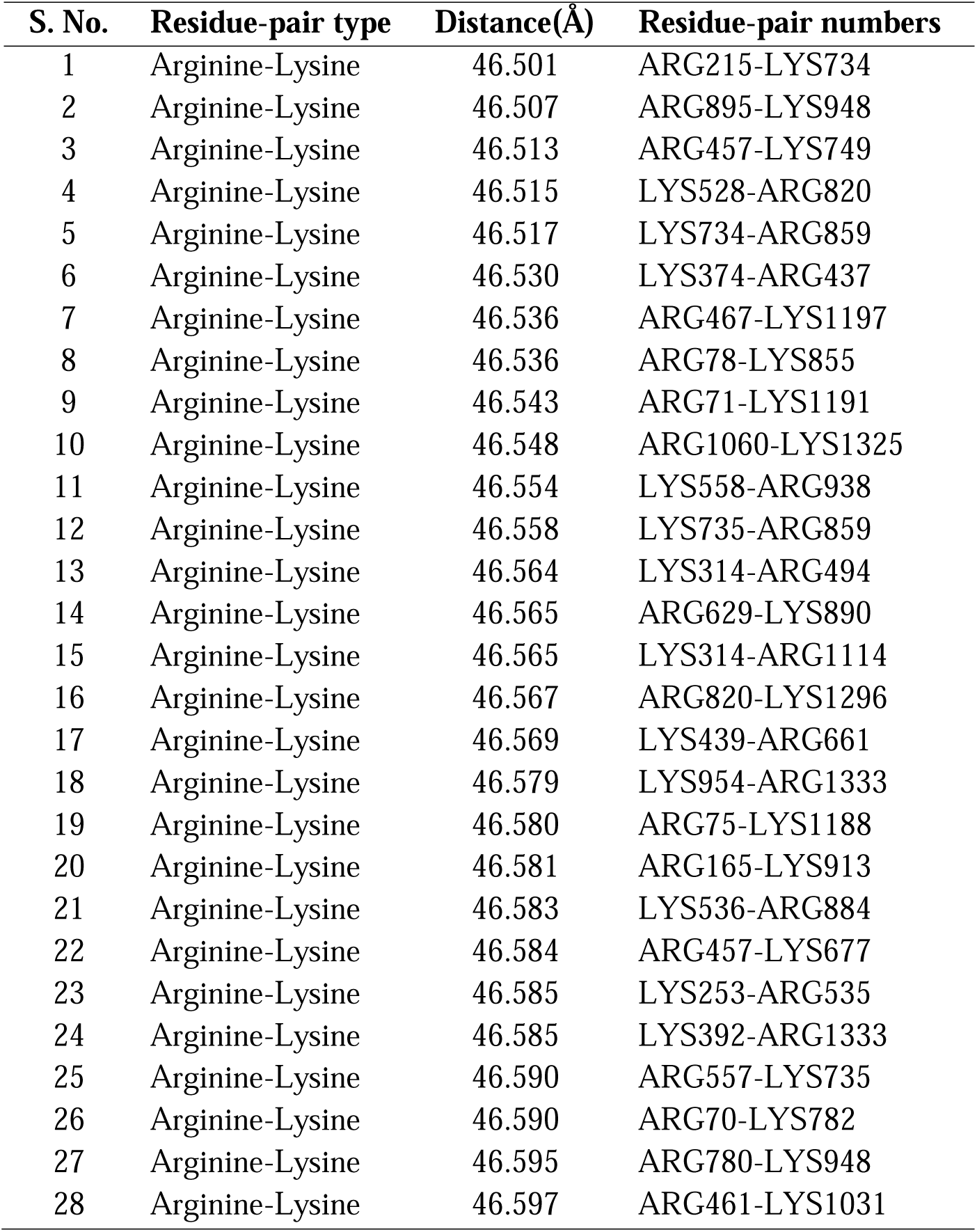
The 28 Lysine-Arginine pairs within top bit of SpCas9 [PDB ID: 5F9R] obtained from second round FS.

We mapped the 28 residue pairs of Table 8 onto the crystal structure of SpCas9 (PDB ID: 5F9R) to visualize their spatial distribution within the protein structure. As shown in Figure 5, these Lysine-Arginine residue pairs are distributed across various key domains of SpCas9, including RuvC, HNH, Rec, and the BH. Notably, the distances between residue pairs spanning different domains suggest their role in maintaining the overall structural integrity of the protein. Interestingly, a significant proportion of these Lysine-Arginine pairs are located near the sgRNA-DNA heteroduplex, where their lysine and arginine residues interact with the negatively charged DNA. This interaction likely enhances the localized stability of the protein in this region, which is critical for its function. Overall, these findings indicate that the identified Lysine-Arginine pairs play dual roles in contributing to both the global stability (spanning multiple domains) and the localized stability (around the sgRNA-DNA binding interface) of the Cas9 structure.

### 3.4. Functional relevance of the identified Lys-Arg Allosteric Network in Cas9

To further investigate the functional significance of the 28 Lysine-Arginine (Lys-Arg) interactions identified in our study, we analyzed the 48 unique Lys and Arg residues within these pairs. Many of these residues have been previously implicated in critical Cas9 activities, including PAM recognition, DNA binding, and cleavage.

#### 3.4.1 Bridge Helix (BH) Arg residues

Among our identified residues, R70, R71, R75, and R78 in the arginine-rich BH form multiple salt bridges with the seed region of the DNA. These interactions facilitate the base pairing of tDNA with sgRNA, stabilizing the R-loop and promoting efficient DNA binding. This structural stabilization plays a critical role in Cas9 function by ensuring precise alignment of tDNA with gRNA^20,63,64^. Experimental evidence further supports the allosteric roles of these BH residues. Nishimasu et al.^20^ demonstrated that mutating R70, R71, R75, or R78 to alanine significantly reduced DNA cleavage activity, emphasizing their importance in seed recognition and cleavage efficiency. Bratovic et al.^63^ mutated multiple of these BH residues and observed reduced cleavage rates across all developed mutants. Notably, Cas9_R71A showed significant DNA binding deficiencies, while Cas9_R70A and Cas9_R74A exhibited increased RNA binding constants, confirming their roles in gRNA binding and R-loop stabilization. Considering these BH residues are well-established allosteric regulators in Cas9, the fact that our Lys-Arg allosteric network analysis independently identified key BH residues, such as R70, R71, R75, and R78, reinforces the validity of our approach and confirms that these interactions are integral to Cas9’s allosteric regulation.

#### 3.4.2 Specificity-related Lysine and Arginine residues

To assess the impact of our identified residues on SpCas9 specificity and off-target activity, we examined the location and function of these residues across different SpCas9 domains. Several residues were directly linked to specificity-enhancing mutations in previous studies. These include BH residues R7 and R78, discussed earlier. Bratovic et al.^63^ analyzed the Cas9_R71A and Cas9_R78A BH mutants to investigate the role of BH arginines in Cas9’s mismatch sensitivity. Their study identified R71 and R78 as key mismatch-sensing residues, with R78 playing a particularly crucial role in recognizing mismatches at target position 4. Structurally, R78 directly interacts with sgRNA nucleotides that base pair with target DNA positions 3-4, highlighting its role in fine-tuning Cas9 specificity.

Likewise, K855 is a key allosteric regulator in SpCas9, facilitating signal transmission between the REC and NUC domains to modulate DNA binding^27,28,65,66^. Molecular dynamics (MD) simulations have demonstrated that this allosteric signaling is mediated by a network of electrostatic interactions between positively charged residues such as K855, K810, and K848, and the negatively charged DNA backbone^30,33,67,68^. Disrupting these interactions, particularly through the K855A mutation in the eSpCas9(1.0) variant, has been shown to impair allosteric communication, leading to a reduction in off-target effects^69^. Given that both R78 and K855 are independently recognized as critical allosteric residues, their identification in our Lys-Arg allosteric network further validates our approach and underscores their importance in Cas9’s regulatory dynamics and specificity.

Several residues within our Lys-Arg allosteric network play essential roles in sgRNA recognition, PAM interaction, and DNA cleavage regulation. R165, located in the REC1 domain, interacts with the backbone phosphate groups of sgRNA, facilitating its recognition. Experimental evidence from Nishimasu et al.^20^ demonstrated that mutating R165 to alanine significantly reduced cleavage activity, underscoring its critical role in SpCas9 function. Similarly, R1333, a key residue in the PAM-Interacting (PI) domain, is crucial for PAM recognition^70^. It forms base-specific hydrogen bonds with dG2 and dG3 nucleobases (5′-NGG-3′) on the non-target DNA strand, and its R1333A mutation drastically reduced target-DNA binding, confirming its importance in sequence specificity. Another PI domain residue, R1114, contributes to Cas9-DNA stabilization by forming multiple ionic and hydrogen bonds with the phosphate backbone of the non-target DNA (ntDNA) strand^71^. Additionally, K890, located in the HNH domain, is among the three residues mutated to generate the Sniper-Cas9 variant, a high-fidelity SpCas9 engineered to modulate target DNA cleavage and improve specificity^72^. While the exact mechanism of allosteric regulation by K890 remains unclear, its presence in our Lys-Arg network suggests a potential role in fine-tuning Cas9 activity through long-range interactions. Furthermore, several of our 48 residues are reported to be experimentally mutated in popular high-fidelity engineered Cas9 variants, such as eSpCas9(1.1), SpCas9-HF^73^, SpCas9HF4^19,73^, and hscCas9-v1.1(HSC.1.1)^74^ designed rationally to reduce off-target effects and enhance Cas9 specificity. These include residues like R1060; a RuvC III residue which is one of the residues mutated experimentally by Slaymater et al. (R1060A)^66^ to develop eSpCas9 (1.0 and 1.1) variants. The identification of these functionally significant residues within our allosteric network analysis further validates our approach and highlights their potential as targets for engineering high-specificity Cas9 variants with reduced off-target effects.

We further identified several specificity-conferring Lys-Arg pairs from the 10,242 interactions in the first round of feature selection. Table S1 highlights 11 of the 10,242 Lys-Arg pairs identified in our analysis that correspond to known mutation sites used in engineered Cas9 variants designed to enhance specificity and reduce off-target effects. Among them, R765-K1246, previously mutated by Zuo et al.^74^ in the development of the high-fidelity SpCas9 variant HSC1.1, was particularly notable. Their study demonstrated that mutations across multiple Cas9 domains reduced off-target activity by weakening interactions with both target and non-target DNA, thereby enhancing specificity. Along with R1060, residues such as K810, K848, and K1003 located in the HNH and RuvC domains are also mutated to design variants eSpCas9(1.0) (K810A/K1003A/R1060) and eSpCas9(1.1) (K848A/K1003A/R1060) respectively. Interestingly, all three pairs, K810-R1060 (54.249Å), K1003-R1060 (20.649Å), and K848-R1060 (20.649Å), appeared in our top Lysine-Arginine pairs from the first round of feature selection. Moreover, Nishimasu et al.^20^ showed that mutating K163, along with R63, R69, R75, and R78, to alanine significantly impaired cleavage, highlighting the critical role of these lysine-arginine interactions in Cas9’s structural and functional integrity. Notably, all four combinations of these lysine-arginine pairs appeared in our top list. This strong alignment with experimentally validated mutations underscores the importance of our Lys-Arg network in defining Cas9 function regulation, including specificity and cleavage activities.

#### 3.4.3 Novel Lys and Arg residues identified in our study

Beyond these previously characterized specificity-enhancing residues, we also identified several novel Lys-Arg interactions that have not yet been reported in Cas9 engineering studies (10242 pairs_Sp_Nme_Cj.xls in SI). Notably, many of these pairs lie within the 5-30Å range near non-target DNA (ntDNA) or the DNA:RNA heteroduplex at the interface of the RuvC, HNH, and PI domains. These interactions may help maintain the heteroduplex structure by forming non-specific electrostatic contacts with the phosphate backbone of DNA. Since these pairs involve positively charged residues interacting with DNA, they may serve as promising targets for neutralization or charge modulation in the design of high-fidelity Cas9 variants with improved specificity. Furthermore, several identified interactions within the L1 and L2 linker regions contain positively charged residues, which could act as hotspots for allosteric modulation. This presents a potential strategy for improving Cas9 catalysis and activity by fine-tuning allosteric communication pathways. However, further experimental validation is necessary to confirm the functional impact of these novel allosteric residues on Cas9 structure and function. The identification of both previously validated and novel specificity-modulating Lys-Arg interactions further validates our approach and provides a structural framework for rational Cas9 engineering.

### 3.5 Molecular Dynamics simulations revealed the distinct roles of Lys-Arg pairs

To investigate whether the identified Lys-Arg residue pairs contribute to functional conformation change of the Cas9•sgRNA•DNA complex, we performed Molecular Dynamics (MD) simulations on the wild-type (WT) SpCas9 system. The simulations tracked distance fluctuations among 28 key residue pairs over a 50–150 ns window. By analyzing these trajectories, we aimed to determine whether these interactions function as allosteric relays, facilitating the coordinated structural changes necessary for DNA recognition and cleavage. Interestingly, while all 28 residue pairs were approximately 46.5 Å apart in the crystal structure (PDB ID 5F9R), their behavior during MD simulations diverged into distinct stabilization trends. These differences likely stem from variations in local structural environments, domain mobility, and their specific roles in allosteric regulation, leading to their classification into three hierarchical groups, as shown in Figure 4.

**Figure 4.**
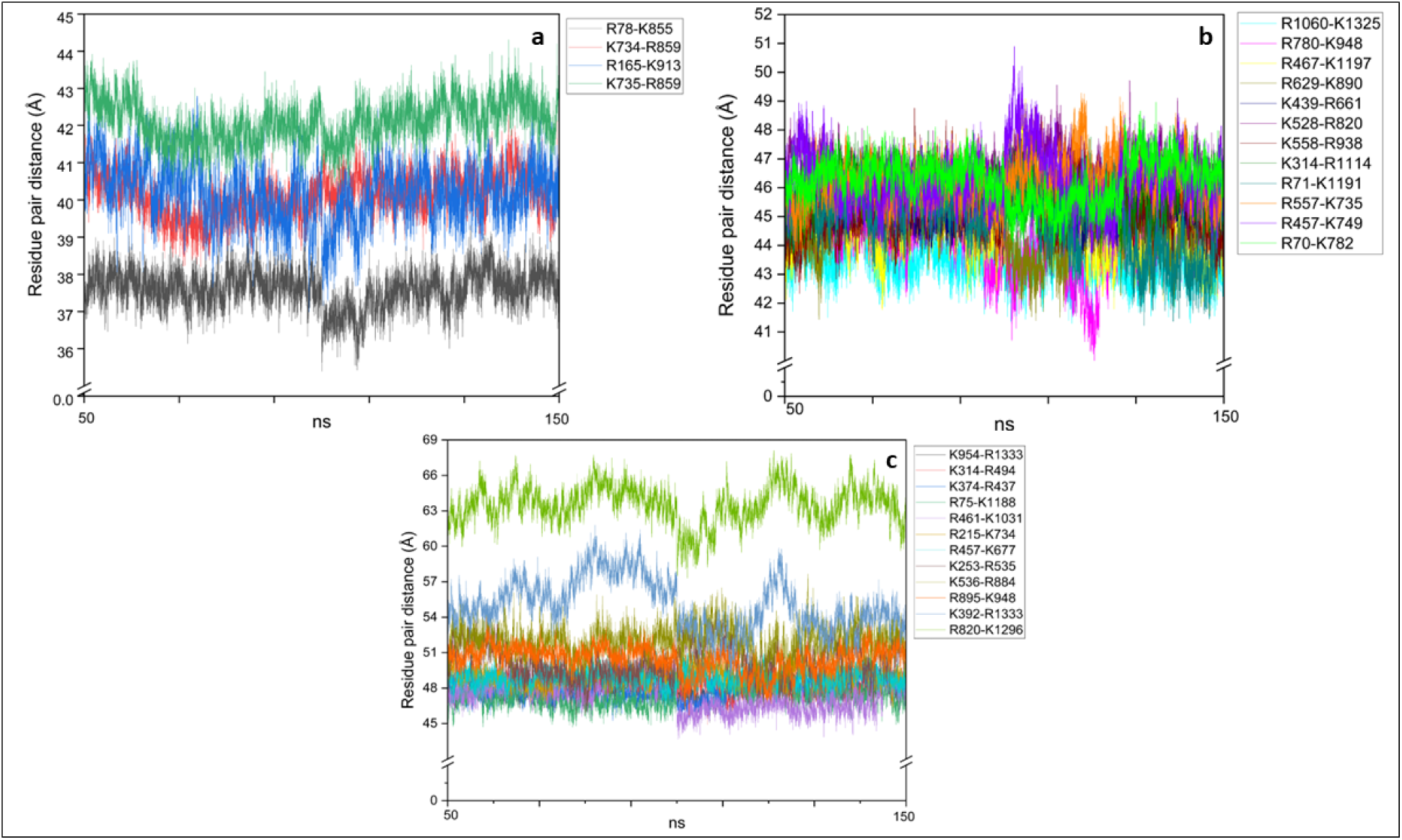
Molecular Dynamics trajectory analysis of the 28 lysine-arginine residue pairs within WT SpCas9 systems. Trajectories representing change in distances of the 28 lysine-arginine residue pairs within the WT SpCas9 system [PDB ID: 5F9R] during a 50-150 ns frame MD simulation.

Figure 4a highlights Group 1 residue pairs, including R78-K855, K734-R859, R165-K913, and K735-R859, which exhibited significant contraction from 46.5 Å in the crystal structure to a stabilized range of 37.0–43.0 Å during simulation. The reduction in distance suggests that these interactions function as allosteric hubs, tightening upon SpCas9 dynamics to facilitate signal transmission between catalytic and DNA-binding domains. For example, the contraction of R78-K855, which bridges the bridge helix (BH) and HNH domains, supports its role as a key allosteric switch regulating cleavage activation. Similarly, R165-K913 and K734-R859, linking the REC and RuvC domains, may help coordinate the conformational changes necessary for target DNA recognition and strand separation.

Group 2 residue pairs, shown in Figure 4b, exhibited smaller distance adjustments, stabilizing within 43.1–46.7Å. These interactions likely act as flexible allosteric relays, maintaining domain connectivity while allowing for controlled structural adjustments. The ability of these pairs to stabilize within a narrow distance range suggests that they contribute to fine-tuning Cas9’s response to DNA binding, ensuring that conformational changes occur in a regulated manner. This behavior aligns with previous findings that allosteric networks within Cas9 must balance stability and flexibility to achieve high specificity without compromising catalytic efficiency.

Figure 4c depicts Group 3 interactions, which stabilized at greater distances (48.0–62.7 Å), with some exhibiting fluctuations throughout the simulation. These residue pairs likely contribute to long-range allosteric regulation, playing a role in maintaining Cas9 in an activation-ready state while allowing for structural adaptability. For instance, interactions such as K954-R1333, which spans the PAM-interacting (PI) domain, may facilitate Cas9’s transition between its DNA-binding and cleavage-competent conformations. Other pairs, such as K536-R884 and K253-R535, show dynamic behavior, suggesting that they provide structural flexibility necessary for Cas9 to accommodate target DNA sequences while maintaining catalytic precision.

The fact that all 28 residue pairs started at the same initial 46.5 Å distance but followed different stabilization trajectories underscores a hierarchical model of allosteric regulation within Cas9. Group 1 interactions act as core allosteric switches, undergoing tightening to initiate conformational transitions. Group 2 pairs function as structural relays, allowing for controlled flexibility between functional domains. Group 3 interactions provide long-range stability, ensuring that Cas9 remains adaptable while preserving its overall structural integrity. The different behaviors observed during MD simulations may be driven by variations in local domain flexibility, interactions with DNA, and their roles in facilitating specific conformational transitions. These findings highlight the importance of long-range residue interactions in modulating Cas9 function.

### 3.7 MD simulations reveal mutation-induced perturbation of the Lys-Arg network and SpCas9 specificity modulation

To explore how mutations perturb SpCas9’s Lys-Arg network and influence specificity, we performed MD simulations on wild-type (WT) SpCas9 and two mutant systems: R78A-K855A (M1) and R765A-K1246A (M2). These mutations were selected to target key long-range Lys-Arg residue interactions identified in our previous analyses. The M1 mutation (R78A-K855A) was chosen due to its location in the bridge helix (BH) and HNH domain, regions involved in inter-domain communication and catalytic activation, and well-established allosteric roles. This pair was also among the 28 residue pairs identified in the second round of FS, reinforcing its potential role in allosteric signaling. The M2 mutation (R765A-K1246A) was identified in our first round of feature selection and was previously reported by Zuo et al.^74^ to enhance specificity, suggesting its involvement in directly modulating DNA interactions. Structurally, R78 and K855 are located on opposite sides of the target DNA (t-DNA), while R765 and K1246 are positioned on opposite sides of the non-target DNA (nt-DNA) strands of the Cas9•sgRNA•DNA tertiary complex (Figure 5c). Our goal was to determine how these mutations perturb allosteric connectivity across the residue network and ultimately modulate specificity.

**Figure 5.**
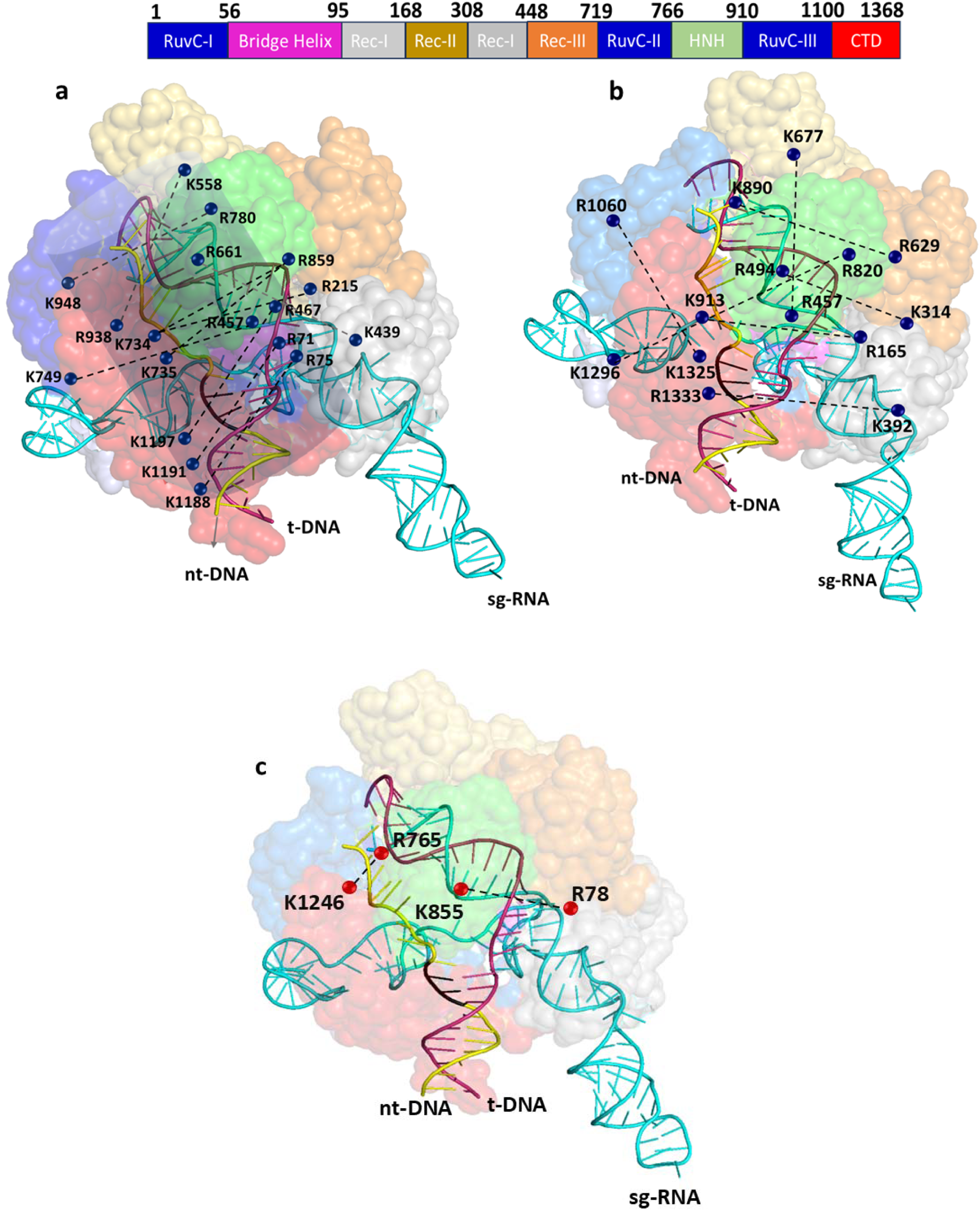
Location of the 28 Lysine-Arginine pairs in SpCas9 structure [PDB: 5F9R] that showed distance changes upon M1 and M2 mutations. **(a)** Lysine-Arginine pairs showing an increase in distance upon M1 and M2 mutation compared to WT over 300nm simulations **(b)** Lysine-Arginine pairs showing a decrease in distance compared to WT upon M1 and M2 mutation over the 300nm simulations. The channel for allosteric signal transmission between the DNA binding and catalytic domain is shown as a blue barrel. **(c)** Cas9•sgRNA•DNA complex showing the location of the mutated residue pairs in M1(R78-K855) and M2 (R765-K1246).

MD simulations were performed for 300 ns on M1, M2, and WT systems. We first investigated whether mutating the key allosteric residues affected the Lys-Arg network listed in Table 8. Our simulations revealed that both M1 and M2 mutations induced widespread perturbations in the Lys-Arg network, affecting long-range interactions beyond the immediate mutation sites.

Among the 28 long-range residue pairs previously identified, we observed a consistent increase in distance in 10 pairs upon M1 and M2 mutation (Figure 5a, Figure S1). These 10 residue pairs were primarily located on opposite sides of the dsDNA, forming key components of the allosteric signaling pathways that connect the DNA-binding and catalytic domains. In the WT system, these positively charged Lys and Arg residues interact with the negatively charged DNA, forming an “electrostatic valley” that stabilizes Cas9-DNA binding and ensures coordinated domain communication during DNA cleavage. This “electrostatic valley” likely plays a crucial role in maintaining inter-domain connectivity, helping Cas9 properly engage DNA while regulating its conformational transitions.

However, when key allosteric switches (R78-K855 and R765-K1246) were mutated, the electrostatic balance within the valley was disrupted, weakening the structural rigidity of these relay points. Without the stabilizing influence of these residues, the electrostatic network lost cohesion, leading to an expansion in inter-residue distances across multiple domains. This increase suggests that these interactions were dependent on the stability provided by M1 and M2, and without these stabilizing interactions, the network became less constrained and more flexible, affecting the structural integrity of Cas9’s DNA-bound state. The observed expansion highlights the interconnected nature of the allosteric network, where perturbing one set of interactions has cascading effects on long-range Lys-Arg pairs, ultimately influencing Cas9’s ability to maintain its active conformation.

On the other hand, eight residue pairs exhibited decreased distances upon M1 and M2 mutations (Figure 5b, Figure S2), suggesting a compensatory tightening effect within the Lys-Arg network. Unlike the increasing-distance pairs, which are positioned on opposite sides of the dsDNA, many of these contracting pairs are located within the same domain or adjacent domains, where they contribute to intra-domain stability. The decrease in distance among these residue pairs likely reflects an adaptive response to the electrostatic valley perturbation caused by M1 and M2 mutations. In the WT system, the balance of positively charged residues stabilizing the negatively charged DNA maintains an even distribution of structural tension across the Cas9 complex. However, when key allosteric switches (R78-K855 and R765-K1246) were disrupted, this balance was altered, leading to a loss of structural support in certain regions. To compensate, other interactions contracted to reinforce stability, effectively redistributing structural tension within Cas9’s allosteric network. This tightening effect likely helps preserve critical structural features of Cas9, particularly in domains essential for catalytic activity and specificity. This dynamic adaptation aligns with a hierarchical allosteric regulation model, where some interactions loosen to enable flexibility, while others tighten to reinforce stability. This interplay between expanding and contracting residue pairs highlights the delicate balance within Cas9’s allosteric network, ensuring the enzyme retains both structural integrity and functional adaptability in response to perturbations.

We further investigated whether mutating M1 (R78A-K855A) could enhance SpCas9 specificity by perturbing the electrostatic valley that stabilizes its interactions with DNA. Previous studies have shown that M2 (R765A-K1246A) mutation enhances specificity by reducing Cas9’s interactions with non-target DNA (nt-DNA) in its active state. We hypothesized that M1 mutation could have a similar effect by disrupting allosteric regulation rather than direct DNA contacts.

A key feature of high-fidelity Cas9 variants is their reduced tolerance for mismatches, which prevents cleavage of off-target sequences. Our analysis showed that both M1 and M2 mutations weakened the “electrostatic valley,” which may increase the energetic barrier for Cas9 to engage with ntDNA, making it more selective for fully matched target sequences. This mechanism is reflected in our binding enthalpy calculations (Figure 6a), where M2 significantly weakens nt-DNA binding (∼35%), a direct effect of disrupting nt-DNA contacts. Despite being far from the nt-DNA binding interface, M1 mutation also induces a reduction in binding energy (∼3.3%), suggesting that it achieves a moderate specificity enhancement through allosteric modulation rather than direct DNA interaction.

**Figure 6.**
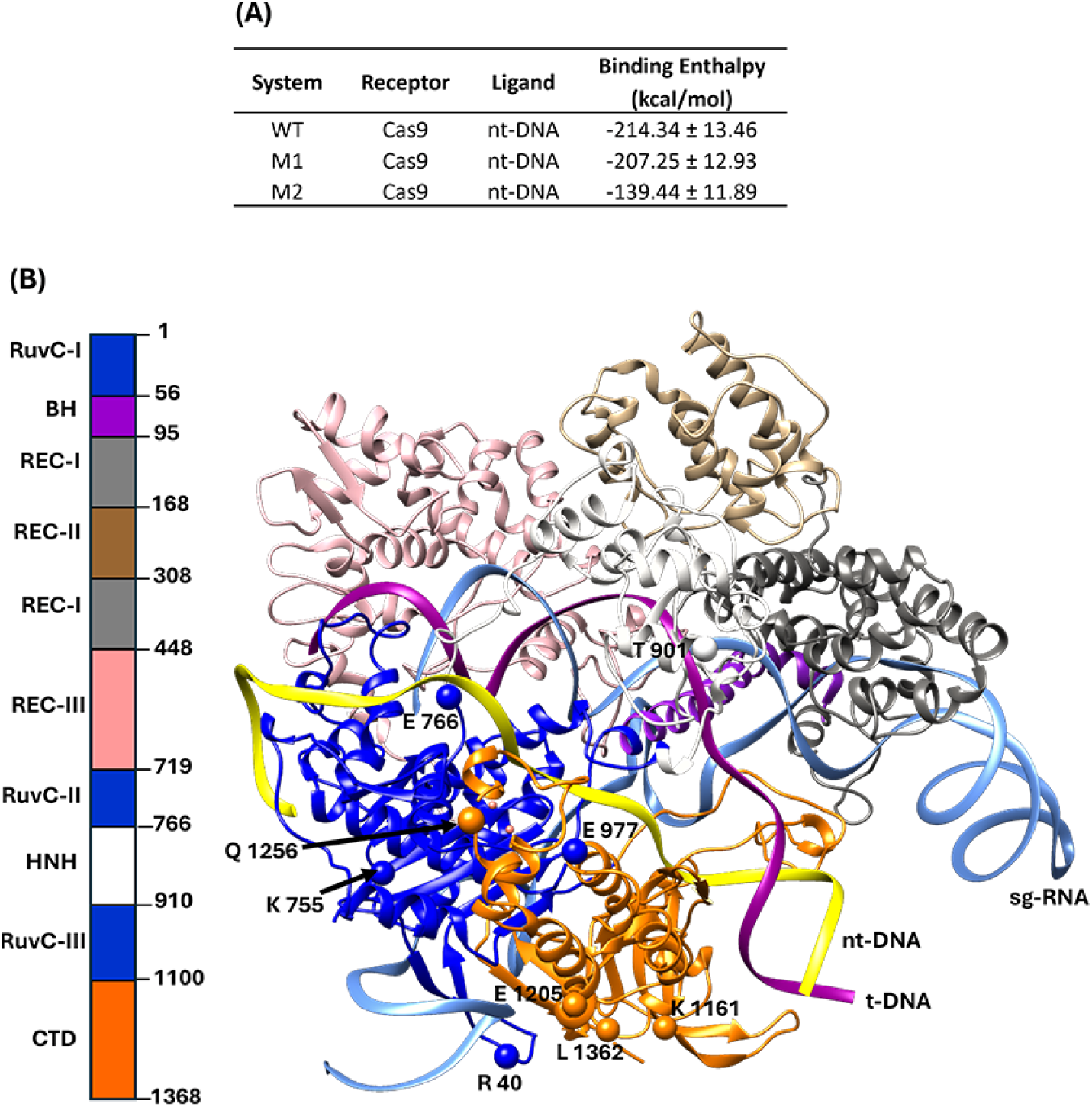
Mutation studies in SpCas9 perturbing the electrostatic valley. **(a)** Average binding enthalpies (kcal mol^−1^) between the SpCas9 (receptor) and the nt-DNA (ligand) for WT, M1 and, M2 calculated based on the MM/GBSA approach on two independent MD simulations. **(b)** SpCas9 residues that show destabilizing effects on nt-DNA binding for both M1 and M2 systems.

The overlay of WT, M1, and M2 Cas9•sgRNA•DNA complexes following MD simulations (Figure S4) revealed a high degree of structural similarity between M1 and M2 (RMSD = 1.71 Å), with both mutations causing significant displacements in the dsDNA and sgRNA duplex regions. This structural resemblance suggests that both mutations exert comparable effects on specificity, albeit through different mechanisms—M2 via direct DNA binding disruption and M1 via allosteric regulation. Figure 6B highlights additional Cas9 residues contributing to nt-DNA destabilization in both M1 and M2 mutants. Notably, none of the Lys-Arg residue pairs from our top distance selections contributed directly to nt-DNA destabilization, reinforcing that Lys-Arg network exerts its specificity-enhancing effects through long-range allosteric perturbations rather than direct interactions.

These findings highlight the potential of combining direct and allosteric modifications to fine-tune Cas9’s specificity. By strategically targeting key long-range residue interactions, it may be possible to develop optimized genome-editing tools that retain high cleavage efficiency while minimizing off-target effects.

## 4. Conclusions

This study introduces a novel structure-based machine learning approach to investigate allosteric regulation in Cas9, offering a new computational strategy for identifying long-range residue interactions that govern enzyme function. While our primary goal is to pinpoint critical residue-pair distances through feature selection, classification serves as a validation step to confirm the significance of these residue pairs identified through feature selection. By employing Cα-Cα inter-residue distances as features, we captured the spatial relationships between amino acids, encoding allosteric networks and inter-residue communications essential for Cas9 stability and specificity. This approach transcends traditional sequence-based methods, leveraging three-dimensional structural features to uncover residue pairs pivotal for interdomain communication and structural integrity. A key innovation in our study is the coarser-to-finer binning strategy combined with a two-stage SHAP-based feature selection (FS), which efficiently reduced the vast feature space of inter-residue distances to a concise set of 28 lysine-arginine pairs without compromising the performance of Random Forest (RF) classification models. The initial coarser binning (5Å) filtered residue pairs by broad distance ranges, while finer binning (0.1Å) refined this to residue pairs within specific intervals (45–65Å), uncovering those crucial for allosteric regulation and stability. This approach first identified 10,242 residue pairs, later refined to 28 significant lysine-arginine pairs in the second stage of FS.

By applying ML-driven feature selection to Cas9’s structural landscape, we uncovered a previously uncharacterized Lys-Arg allosteric network, which plays a fundamental role in stabilizing Cas9’s DNA-bound conformation. MD simulations revealed a hierarchical allosteric regulation model, where different subsets of Lys-Arg redistribute structural tension, facilitating conformational transitions essential for its function. Mutational analyses of R78A-K855A (M1) and R765A-K1246A (M2) revealed that these Lys-Arg networks form an “electrostatic valley” along the Cas9-ntDNA interface, where positively charged residues interact with negatively charged DNA to ensure structural integrity. Disrupting this “electrostatic valley,” either through direct or indirect mutations, results in structural destabilization, affecting Cas9’s ability to engage with DNA through non-specific interactions and altering Cas9 specificity.

Overall, our study provides a new framework for studying allostery using ML-driven structural analysis and introduces the electrostatic valley as a key concept in Cas9 regulation. By mapping and perturbing allosteric residues, we offer a rational approach for engineering high-fidelity Cas9 variants, balancing on-target activity with enhanced specificity. Our integrated ML-MD methodology can also be applied to other biological systems, broadening the understanding of allosteric regulation in enzyme function and therapeutic development.

## Acknowledgments

This work is supported by a grant from the National Institute of General Medical Sciences of the National Institutes of Health (R21GM144860).

## Code and Data Availability

The datasets of Cas9 and non-Cas9 proteins are given as supporting information in the “Datasets.xls” file. The global ranking for all the bits in the first round FS is given in supporting information as file “Ranks of first-round bits.xls.” Individual SHAP feature importances of all first-round bits in each of the 15 splits and their average feature importances are also given in supporting information as in the file “Individual SHAP values of first-round bits in 15 runs.xls”. The 10,242 lysine-arginine pairs of SpCas9 and the lysine-arginine pairs in the top bits of NmeCas9 and CjCas9 obtained in the first round FS are given in the file “10242 pairs_Sp_Nme_Cj.xls”.

The python codes use for pre-processing steps, feature selection and RF modeling is give at https://github.com/Sireesiru/Preprocessing-codes-for-Cas-NonCas-classification

## Author Contributions

JL and SW conceived the idea, provided support to develop the workflow, reviewed the manuscript critically, and monitored the entire study. SSM designed the study, developed the ML models, contributed to the result analysis, and drafted the manuscript in coordination with other authors. VMJA performed the MD simulations and computational mutation studies, contributed to trajectory analysis, and drafted the manuscript. CNR collected and cleansed the RCSB-PDB Cas9, AlphaFold-2, and non-Cas9 protein structures and assisted with feature generation tasks. All the authors have reviewed the manuscript critically and given intellectual input, collectively contributing to the study.

