## Supporting tables and figures for "Structure-Based Classification of CRISPR/Cas9 Proteins: A Machine Learning Approach to Elucidating Cas9 Allostery"

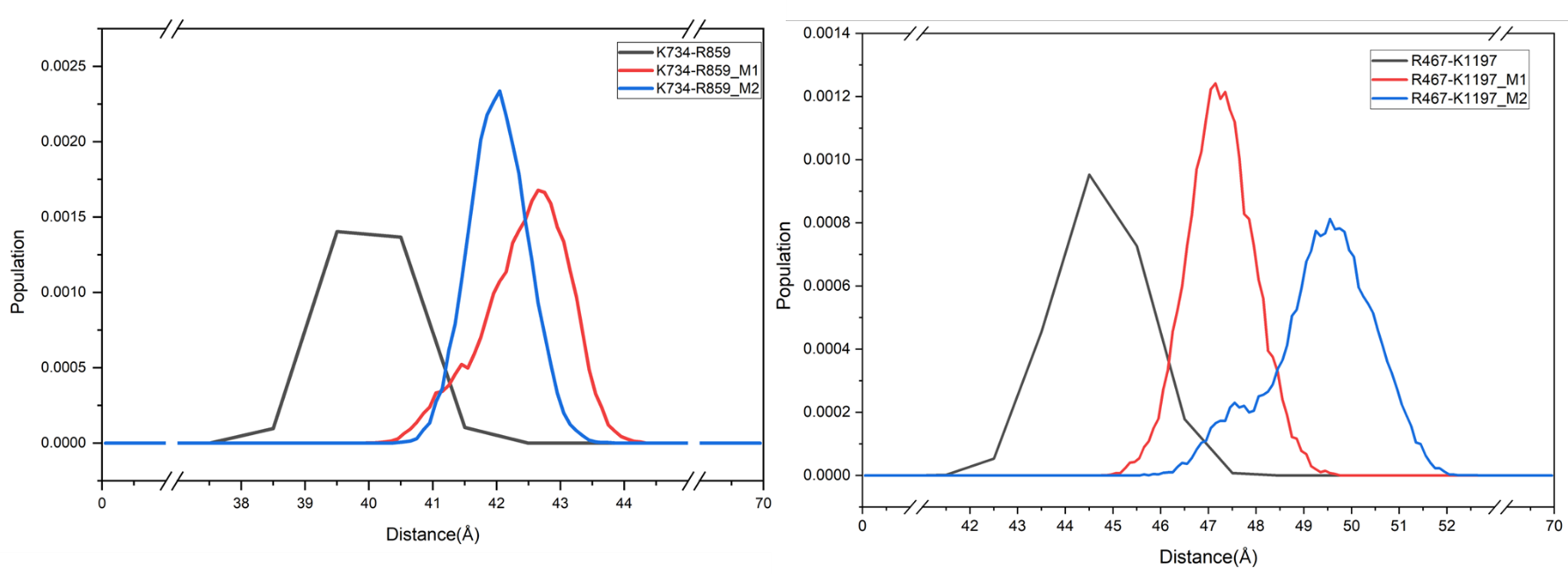
**Figure S1** The 10 pairs showing increasing trend of lysine-arginine distances upon M1 and M2 mutation.

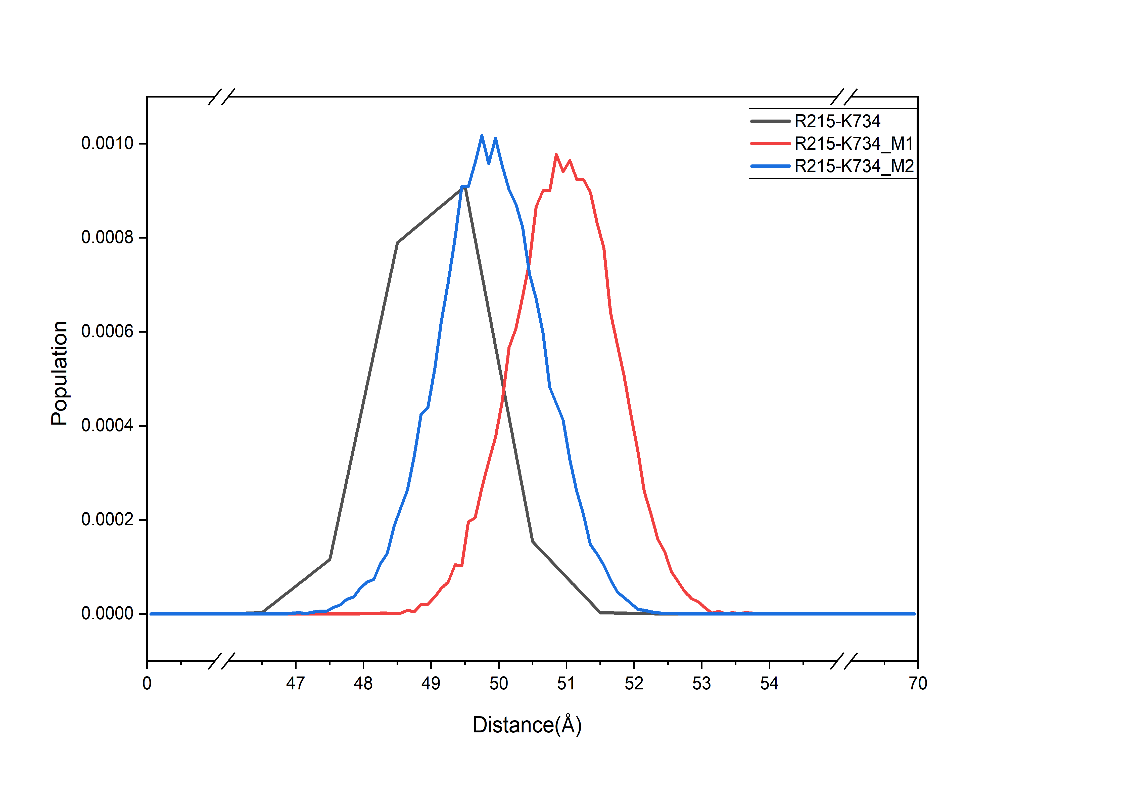

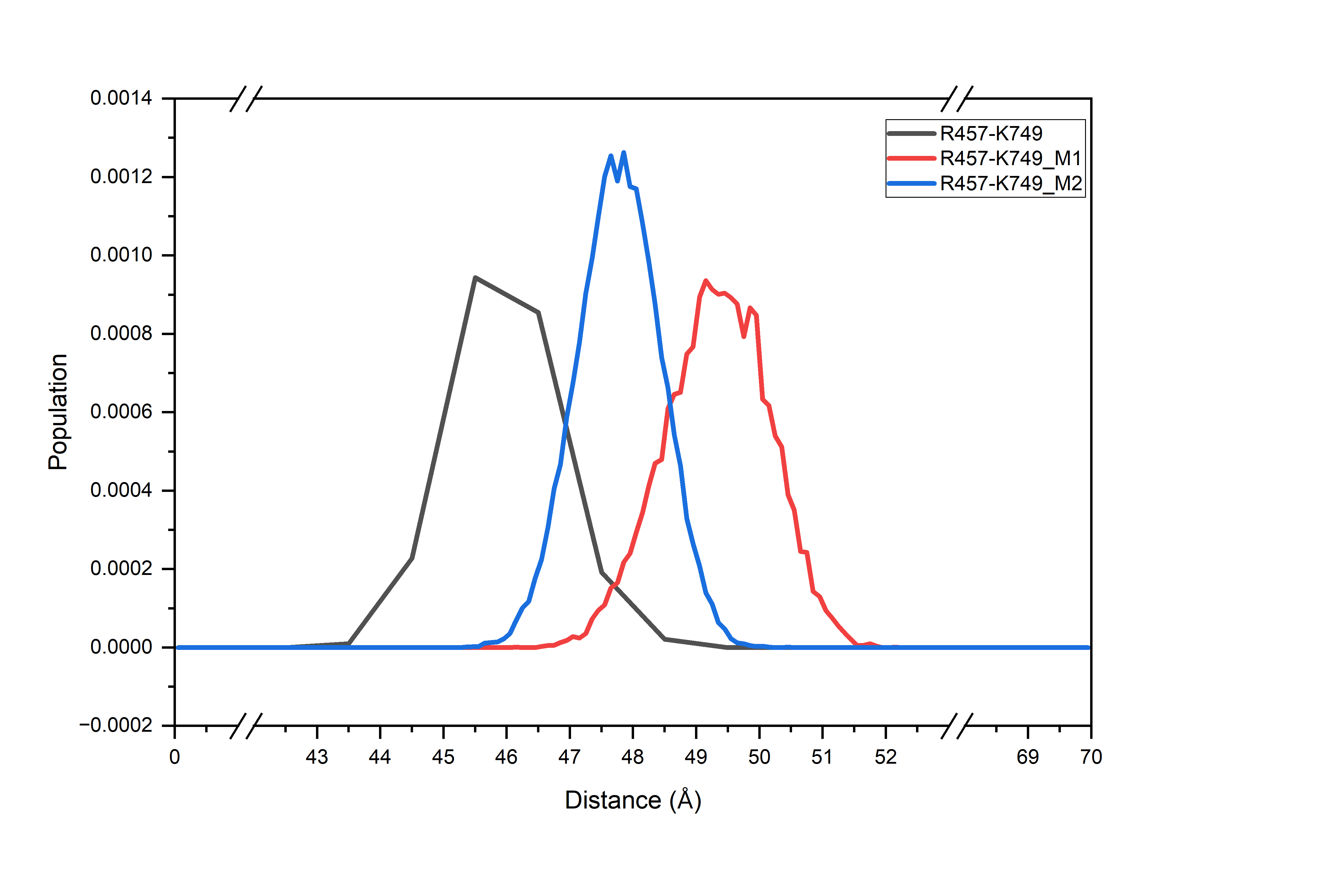

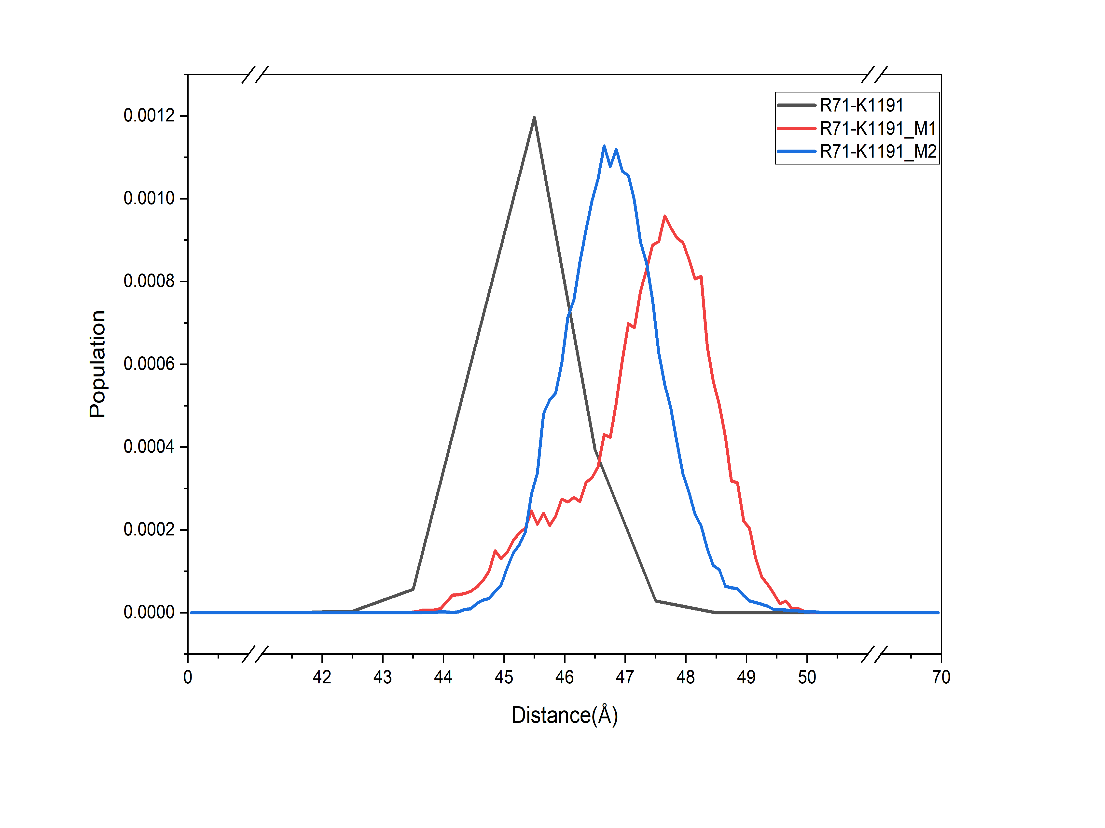

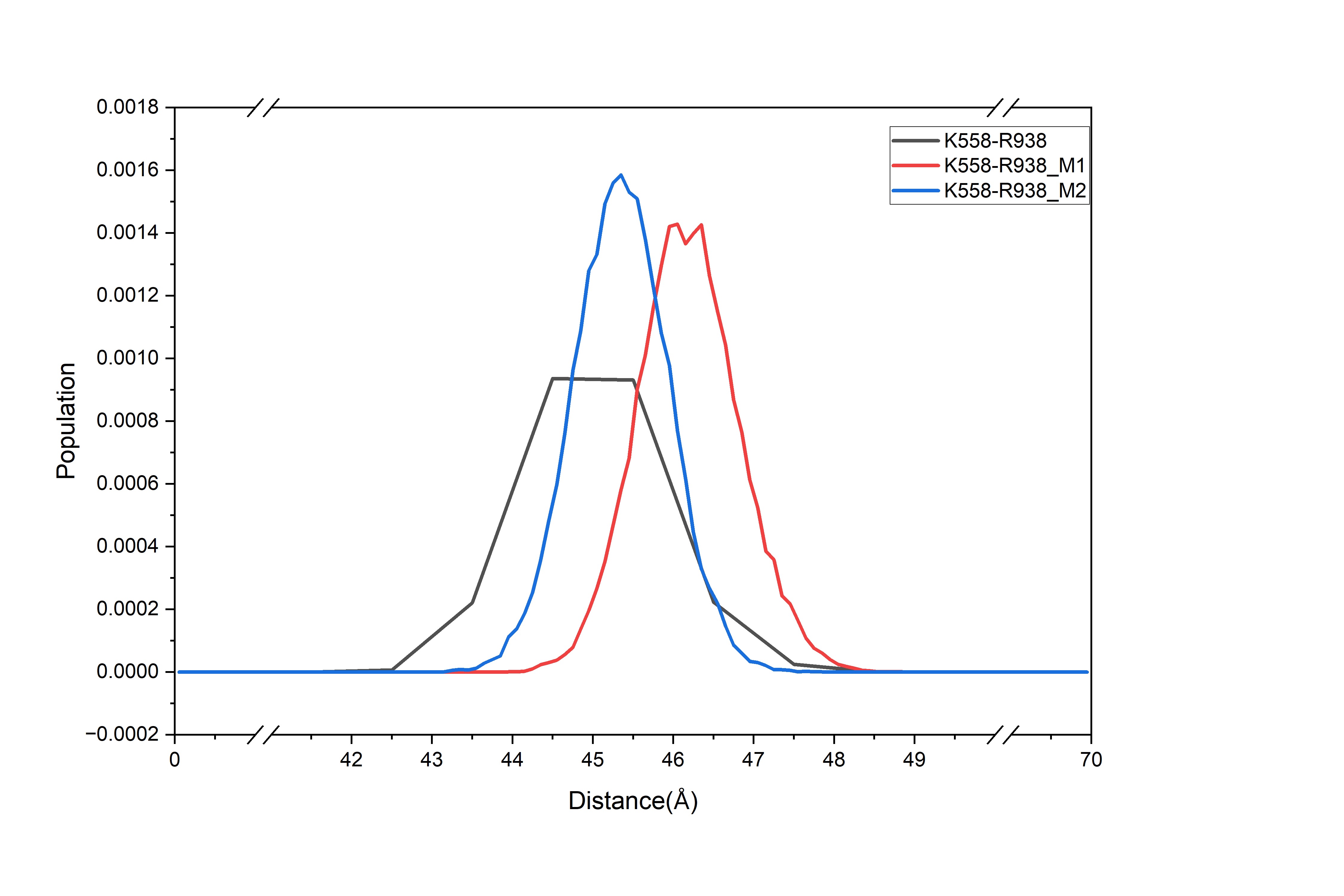

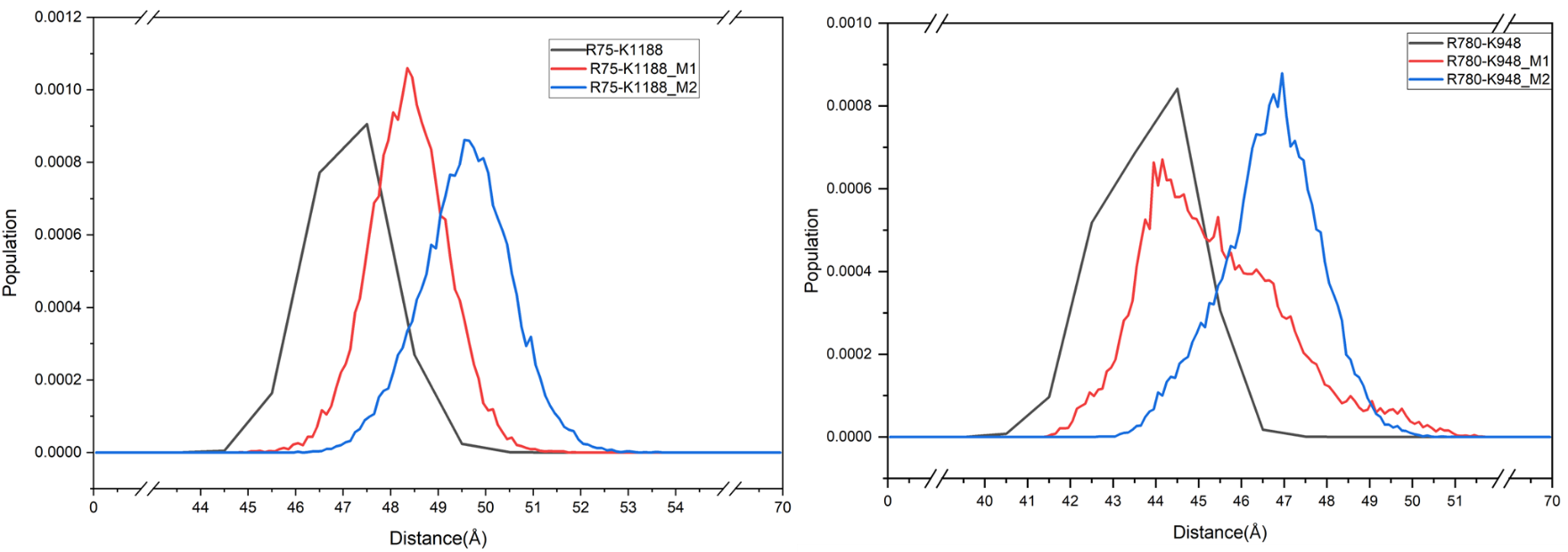

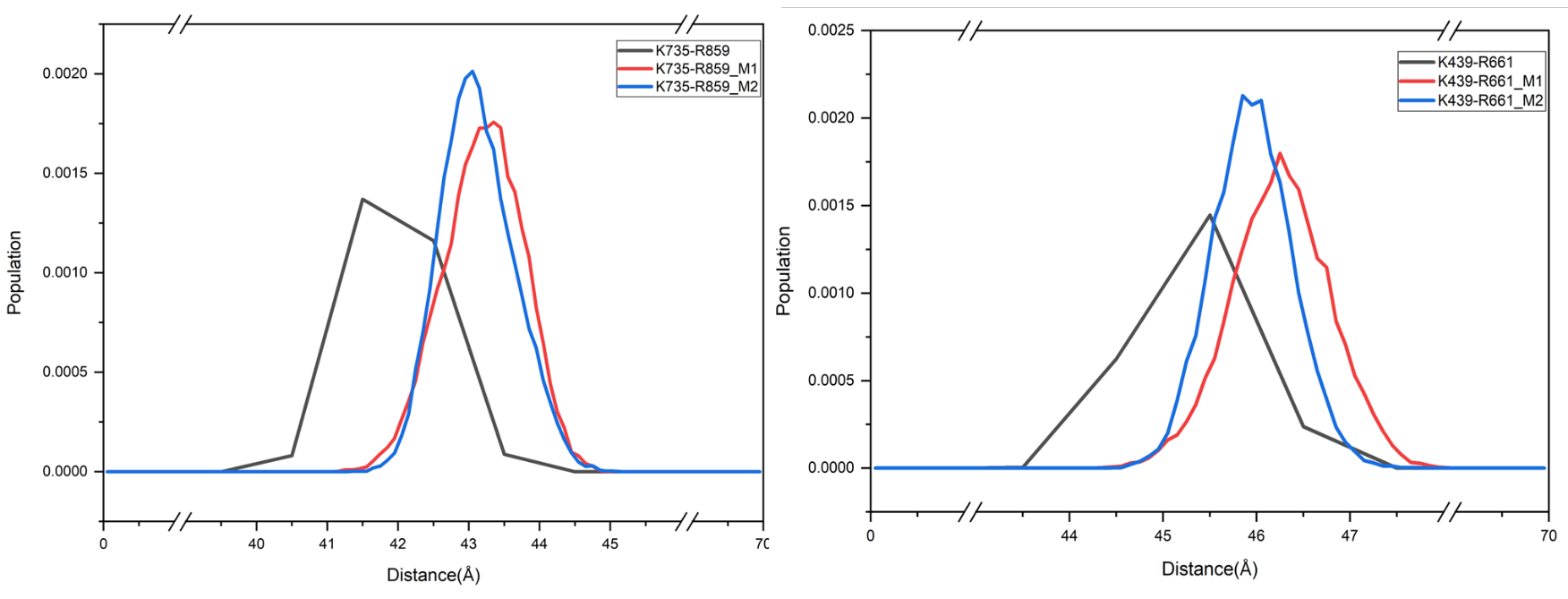

**Figure S2** The 8 pairs showing decreasing lysine-arginine distances
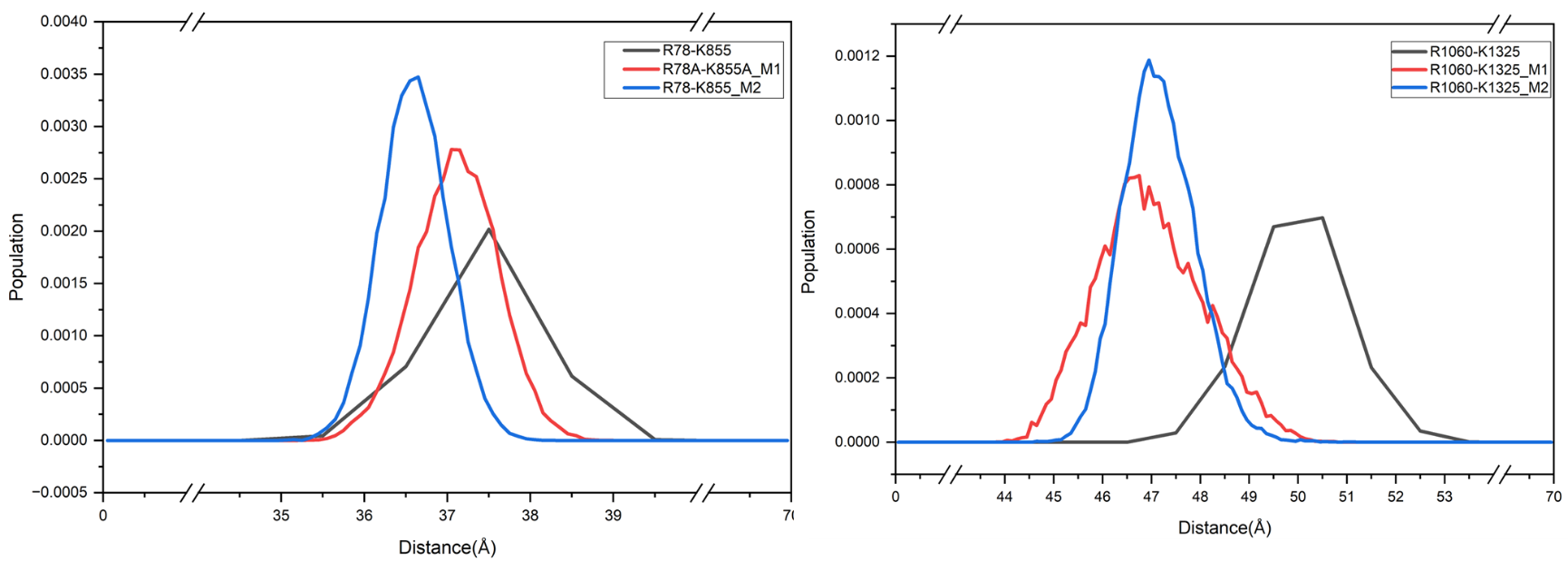
 upon M1 and M2 mutation

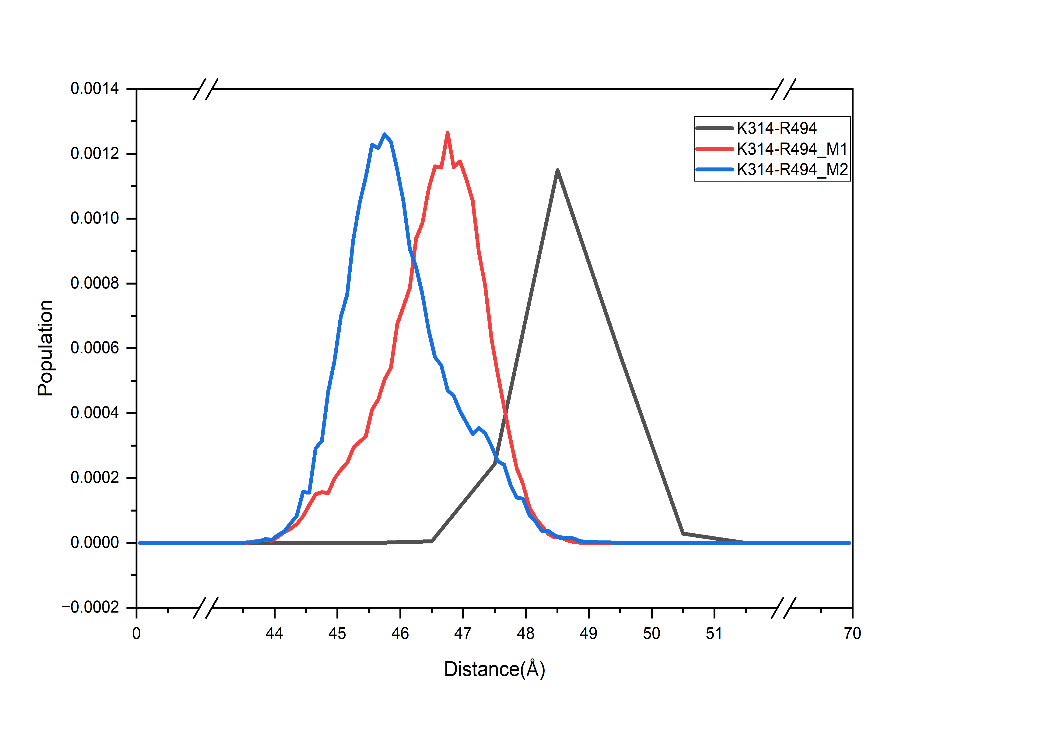

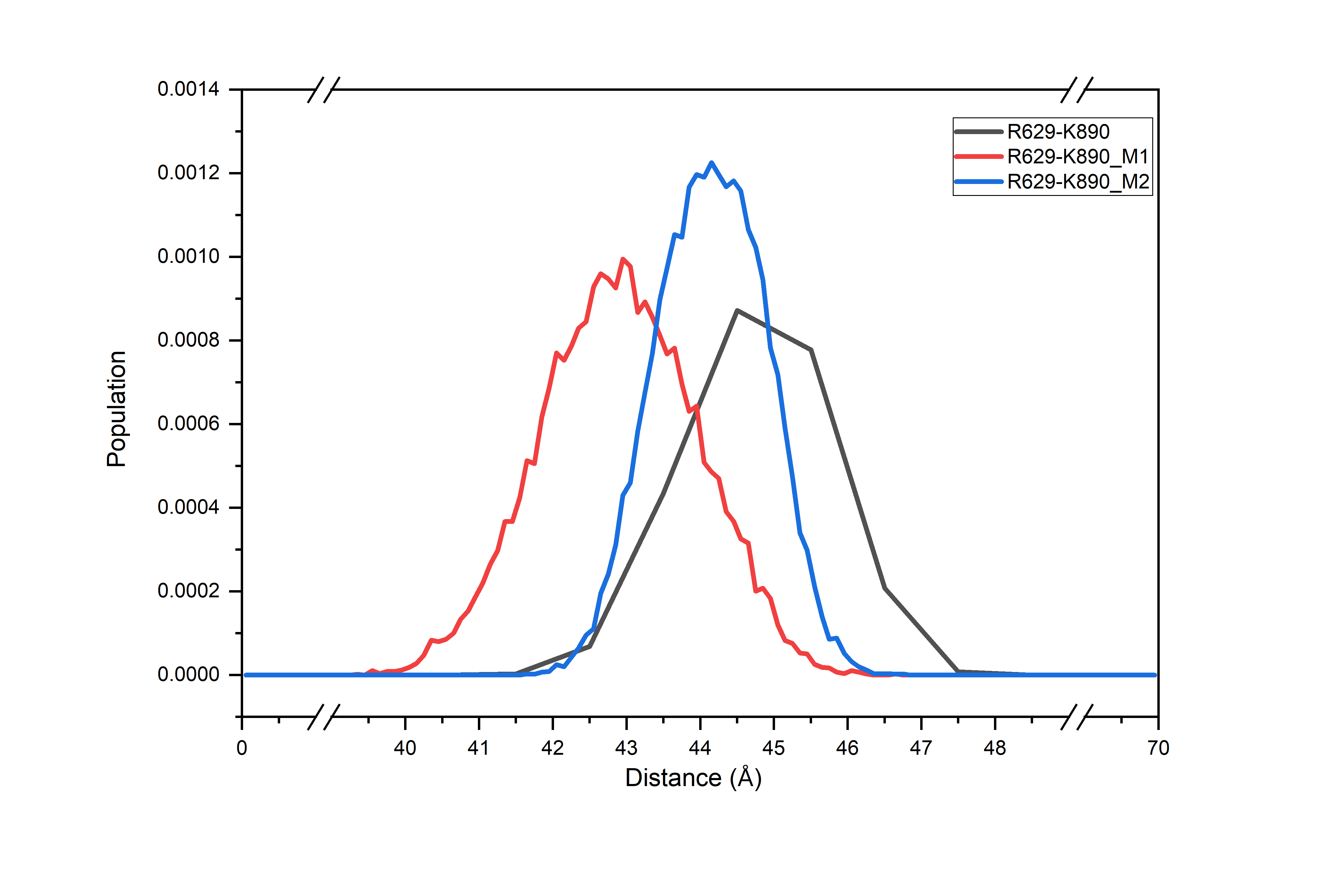

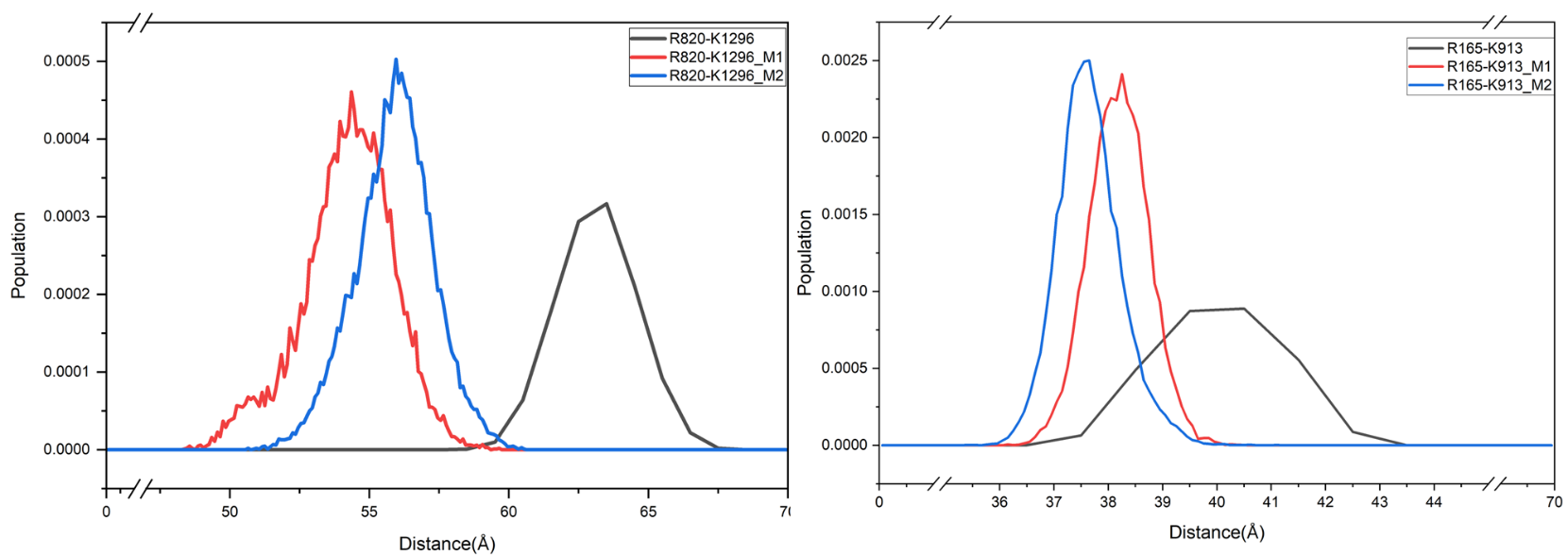

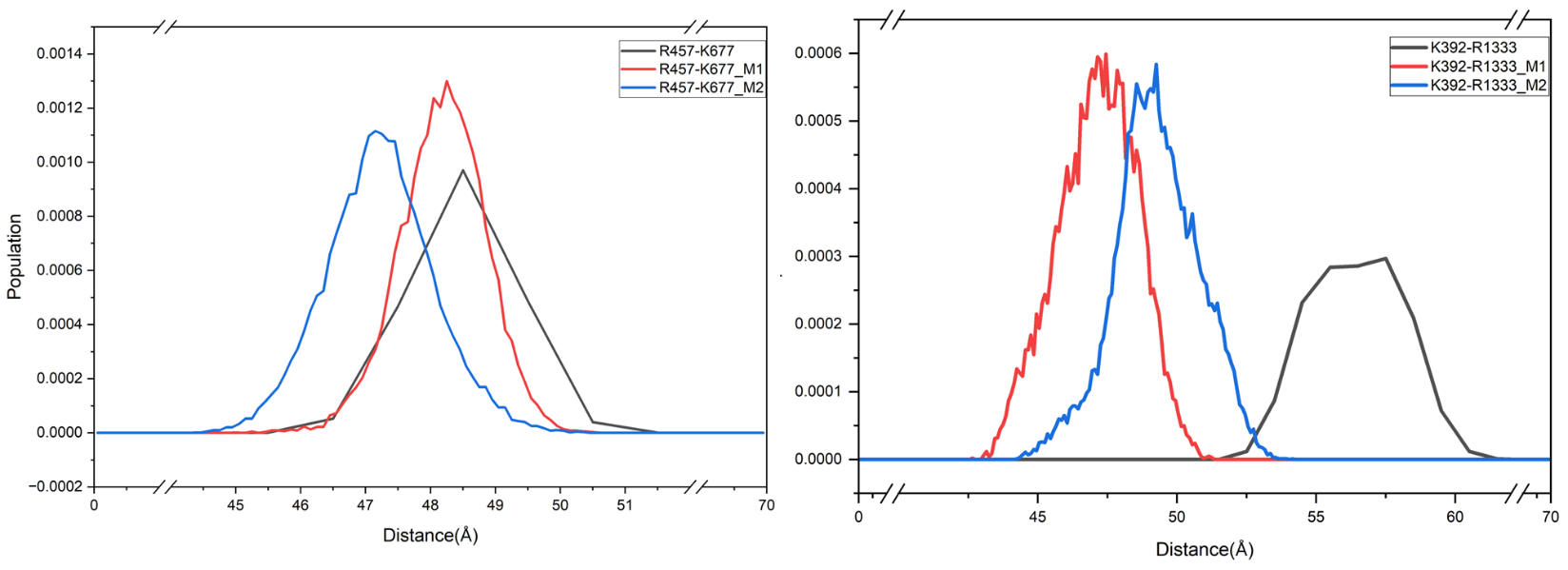

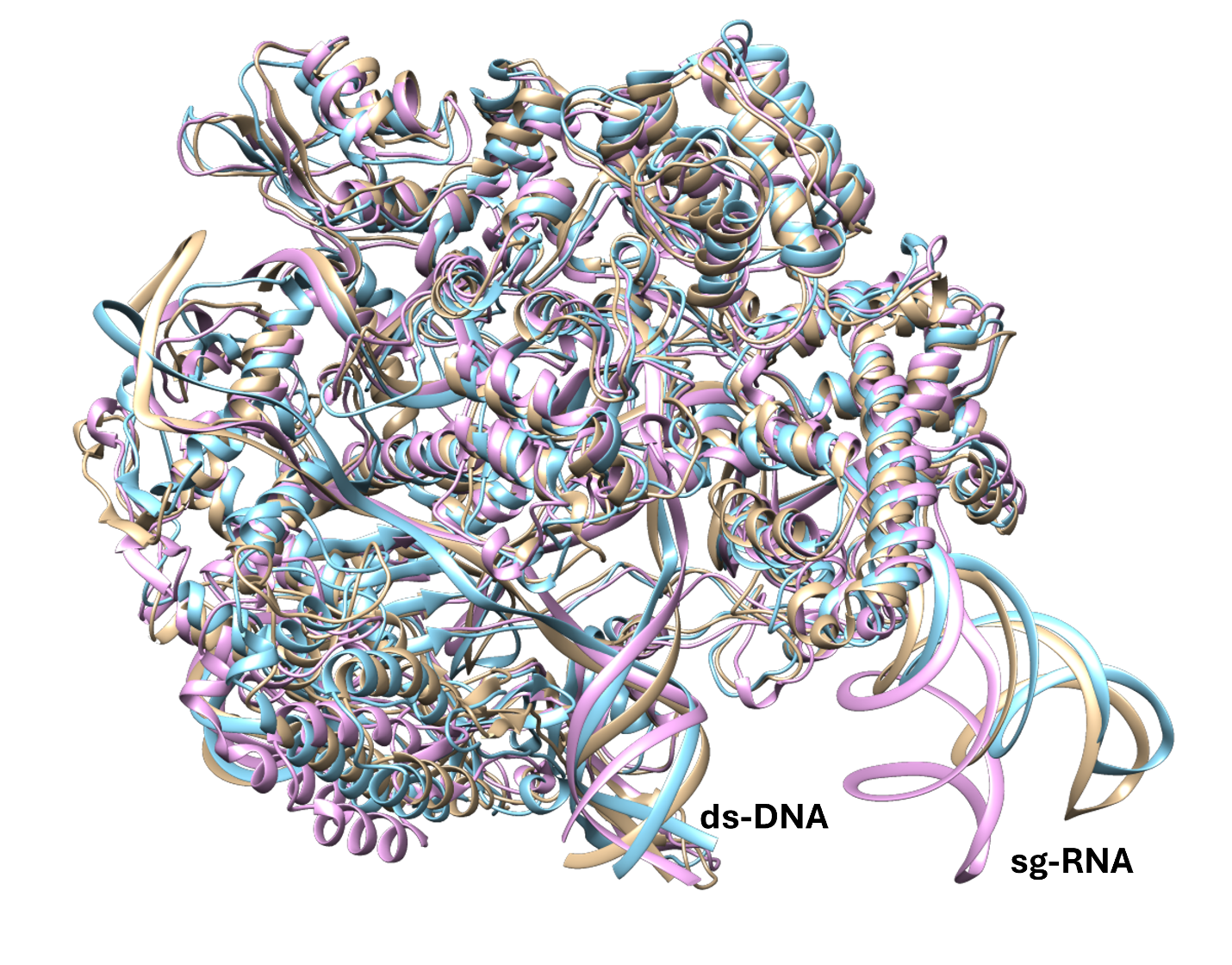

**Figure S3**: An overlay of the most representative tertiary structures (Cas9•sgRNA•DNA) of WT (purple color), M1 (kaki color) and, M2 (blue color) obtained from MD simulations.

**Table S1.** Lysine-arginine pairs from our first round of FS aligning with reported mutations that led to improved specificity or reduced off-target effects in Cas9-derived genomic editors and Cas9 variants.

| **Distance(Å)** | **Pair_num in our study** | **Editor** | **Residues mutated from literature** | **Reference** |
| --- | --- | --- | --- | --- |
| 08.381 | K510-R661 | V3 | N497A, **K510A, R661A** | (Wang et al. 2024)^57^ |
| 20.647 | K1003-R1060 | eSpCas(1.0)  eSpCas(1.1) | K848A, **K1003A, R1060A** | (Slaymaker et al. 2016)^48^ |
| 20.934 | K526-R661 | evoCas9 | **K526E, R661Q** | Casini et al. 2018)^58^ |
| 23.166 | R765-K1246 | Cas9-v1.1 (HSC1.1) | **R765A**, D835A, **K1246A** | Zuo et al. 2022^56^ |
| 25.473 | K848-R1333 | eCas9-SpRY | A61R, **K848A**, K1003A, R1060A, L1111R, D1135L, S1136W, G1218K, E1219Q, N1317R, A1322R, **R1333P**, R1335Q, T1337R | W. Zhang et al. 2021a, b, c^59–61^ |
| 28.963 | K848-R1335 |  | A61R, **K848A**, K1003A, R1060A, L1111R, D1135L, S1136W, G1218K, E1219Q, N1317R, A1322R, **R1333P**,R1335Q, T1337R | W. Zhang et al. 2021a, b, c^59–61^ |
| 33.665 | R661-K848 | HeFSpCas9 | N497A, R661A, Q695A, K848A, Q926A, K1003A, R1060A | (Kulcsár et al. 2017) ^62^ |
| 34.497 | R661-K1003 | Opti-SpCas9 | R661A, K1003H | (Choi et al. 2019)^63^ |
| 44.329 | K848-R1060 | eSpCas (1.1) | **K848A**, K1003A, **R1060A** | Slaymaker et al. 2016^48^ |
| 46.623 | K855-R1060 |  |  | patent/US-10876100-B2 |
| 54.249 | K810-R1060 | eSpCas (1.0) | **K810A**, K1003A, **R1060A** | (Slaymaker et al. 2016)^48^ |

*Boldfaced residues are reported mutations from literature causing increase in specificity or reduced off-target effect. They are the residues that overlap with residues in our identified pairs
